# An open interface system for non-invasive brain-to-brain free communication between naïve human participants

**DOI:** 10.1101/488825

**Authors:** Angela I. Renton, Jason B. Mattingley, David R. Painter

## Abstract

Free and open communication is fundamental to modern life. Brain-computer interfaces (BCIs), which translate measurements of the user’s brain activity into computer commands, present emerging forms of hands-free communication. BCI communication systems have long been used in clinical settings for patients with paralysis and other motor disorders, and yet have not been implemented for free communication between healthy, BCI-naïve users. Here, in two studies, we developed and validated a high-performance non-invasive BCI communication system, and examined its feasibility for communication during free word association and unprompted free conversation. Our system, focusing on usability for free communication, produced information transfer rates sufficient and practical for free association and brain-to-brain conversation (~5.7 words/minute). Our findings suggest that performance appraisals for BCI systems should incorporate the free communication scenarios for which they are ultimately intended. To facilitate free and open communication in healthy users and patients, we have made our source code and data open access.

## Introduction

### Overview

Free and open communication is fundamental to modern life, scientific enterprise and democratic discourse. The rising prevalence of technologies such as virtual/augmented reality ^[1–4]^ and artificial intelligence ^[5; 6]^ has created new opportunities for hands-free communication and control. Brain-computer interfaces (BCI), which translate measurements of the user’s brain activity into computer commands to control external devices ^[7–11]^, present emerging forms of hands-free communication. BCI spellers are virtual keyboards that decode brain activity patterns allowing users to select characters in sequence to spell words and, ultimately, freely communicate ^[12]^. BCI keyboards mimic manual keyboards, which extend the user by allowing them to physically manifest their real-time thoughts, interface with the internet and communicate remotely. BCI communication systems, including spellers, have long been used in clinical settings to facilitate communication in cases of quadriplegia, anarthria and amyotrophic lateral sclerosis ^[13–15]^. These systems are often developed using electroencephalography (EEG), which allows for portable, flexible and affordable devices ^[16]^. BCI has the potential to revolutionise communication, and yet its potential for creating free communication in healthy users is largely unexplored ^[17]^. Here we introduce a new and efficient non-invasive system, using sparse-electrode electroencephalography (EEG), that allows free communication between individuals based on real-time brain activity decoding.

Remarkable progress had been made in the development of signal processing and classification algorithms to decode the brain activity underlying BCI control commands ^[12; 18–21]^, including faster and more accurate classification of neural activity evoked by flickering virtual keyboards ^[6; 12; 21; 22]^. For communication, the original and most prevalent BCI speller uses the P300, an event-related potential (ERP) component modulated by attention. P300 spellers require multiple iterations for single character selection, and it may take minutes to type a single word ^[23–29]^ when not combined with predictive text ^[30]^. Spellers based on the steady-state visual evoked potential (SSVEP), which rely on a combination of gaze shifting and attention-related entrainment of visual cortical neurons to flicker frequency, allow higher information transfer rates (ITRs) due to increased signal-to-noise ratio (SNR) of SSVEPs relative to ERPs ^[22; 31–35]^. For instance, Chen et al. ^[21]^ and Nakanishi et al. ^[36]^ developed SSVEP-based spellers with information transfer rates (ITRs) of up to ~325 bits min^−1^ (bpm) for cued spelling, with a mean classification accuracy of over 90%. These ITRs are the fastest to date, with other state-of-the-art spellers averaging ~146 bpm ^[34; 37–41]^*. This impressive improvement suggests that BCI spellers may become a viable option for hands-free communication outside of clinical settings.

Despite advances in signal processing and classification, BCI spellers have rarely been explored for free communication in healthy individuals. Free communication involves translating momentary thoughts to text, continuously and in real time. However, current tests of “free” spelling often involve users repeating a small number of phrases provided by the experimenter, either from memory or with assistance from salient cues ^[21; 36]^. While algorithms may allow ultra-high ITRs on cued tests, it is not clear that naïve users can cope with the significant cognitive load associated with BCI operation to freely communicate at these rates. Consider for instance the virtual keyboard developed by Chen et al. ^[21]^, which produced ITRs of ~267 bpm. Their approach was to combine joint frequency/phase modulated flicker with filter-bank canonical correlation analysis (CCA), providing high accuracy for large set sizes (40 keys) and short trial durations (1 s). The reported ITRs, however, likely overestimate free communication speed for several reasons. First, the classification assessment consisted of users BCI typing the phrase “HIGH SPEED BCI” three times, interleaved by one-minute breaks. Testing on a single, cued phrase scarcely resembles free communication, which involves the increased cognitive load of generating thoughts, planning phrases, spelling words, and locating the correct characters, all in real-time. Second, in previous work, ITR was calculated based on a set size of 40 keys. Although all keys were used for template generation, only nine were actually used for typing at test (i.e., “H”, “I”, “G”, “S”, “P”, “E”, “D”, “B”, “C”), violating the preconditions of ITR calculation ^[18; 42; 43]^. Finally, in the study of Chen et al. ^[21]^, the majority of participants were experienced users, having trained on previous BCI systems, as well as the 200 practice trials in which target letters were highlighted by salient cues. The efficacy of such spellers for genuinely free communication in BCI-naïve individuals is therefore yet to be established^[17; 44]^.

### A Non-Invasive Interface for Brain-to-Brain Free Communication

BCI systems seem promising for the development of hands-free communication. We
therefore developed a state-of-the-art filter-bank CCA SSVEP BCI speller (**Figure 1**)^[21; 38]^, and examined its suitability for genuine free communication. In Experiment 1, we tested whether naïve users could maintain rapid typing during prompted free word association ^[45]^. In Experiment 2, we developed a social BCI communication interface, allowing two experienced users to have a free conversation. To facilitate free communication, we introduced eight important changes to previous top-performing virtual keyboards^[21; 38]^.

#### Reductions in character selection time

(1) We changed the keyboard layout from alphanumeric to QWERTY, which is highly familiar to users ^[46]^. (2) We reduced the number of flicker frequencies (from 40) to 28, excluding numbers and punctuation characters, which were deemed superfluous to free communication. (3) We displayed the three last classified characters at the top of every key, allowing users to monitor decoding while entering virtual keystrokes.

#### Increases in classification accuracy

(4) Presenting status characters at the top of each key reduced working memory load as well as the number of saccades required between classification intervals. (5) To reduce potential interference from endogenous alpha oscillations, we used a higher frequency range (10.0 - 15.4 Hz rather than 8.0 - 15.8 Hz). (6) We developed a procedure to calculate the optimal phase shift for the sinusoid templates, thereby tailoring the templates to each individual. (7) We increased the flicker period (from 0.50) to 1.50 s, and the flicker-free interval (from 0.50) to 0.75 s. (8) We developed an ERP method to increase SSVEP SNR, allowing participants with low classification accuracy to potentially use the BCI communication system.

**Figure 1.**
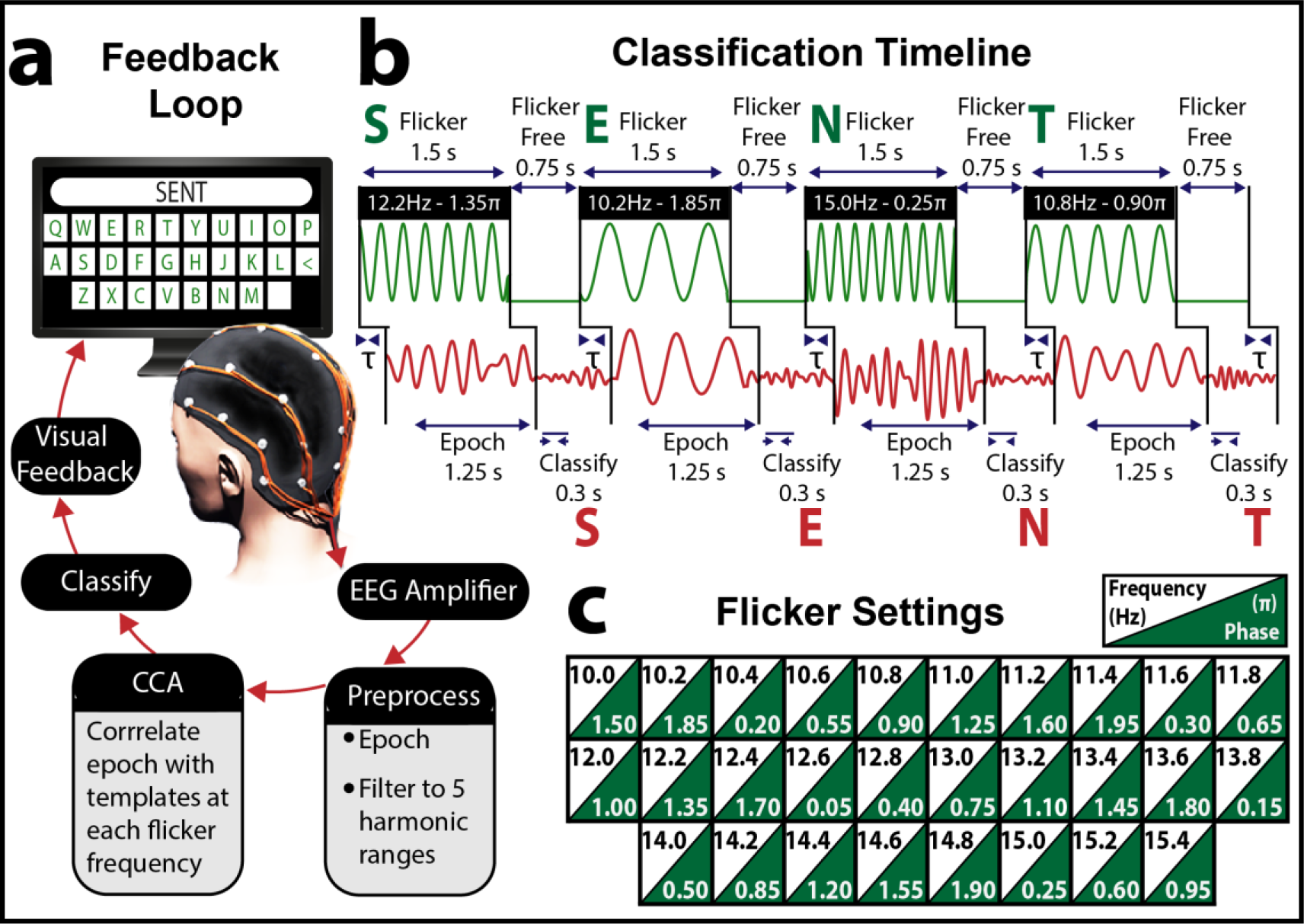
BCI virtual keyboard for free communication. (***a***) Participants operated the real-time feedback loop to freely type words and phrases using their brain activity alone. Participants selected characters in sequence by focusing their attention and fixating their gaze on sinusoidally flickering keys of a virtual QWERTY keyboard on a computer display, which evoked oscillatory SSVEP responses at the corresponding flicker frequency/phase in the EEG. EEG time-locked to flicker was extracted, bandpass filtered to five harmonic ranges and then submitted to a filter-bank CCA with respect to a bank of individualized training templates. The classified frequency was the template most highly correlated with the real-time EEG, with the corresponding character displayed as feedback at the top of each key. Participants were free to select the next character, or to select the backspace key [<] to make a correction. (***b***) Example timeline of visual stimulation and evoked EEG involved in BCI typing of the word “SENT”. Each key flickered at a unique frequency/phase for 1.5 s, followed by a 0.75 s flicker-free period, during which the letter was classified and participants shifted their attention to the next key. Focusing attention on a key potentiated the corresponding SSVEP response, increasing the likelihood that the corresponding letter would be classified. *τ* refers to the SSVEP delay relative to flicker onset, calculated separately for each frequency and harmonic. (***c***) Spatial organisation of the virtual keyboard’s flicker frequencies/phases. Each key flickered at a unique frequency/phase, ranging from 10 Hz/1.5π - 15.4 Hz/0.95π.

## Experiment 1: Results

### QWERTY Classification Assessment

Classification templates derived from cued training (*N* = 20 repetitions/key; **Figure 2*a***) were evaluated by having naïve participants freely BCI type the complete QWERTY sequence three times, without guiding cues other than status characters printed on each key (**Figure 2*b***). This test revealed that the BCI naïve participants could use the communication system with varying degrees of voluntary control, with classification accuracy from 22.62 - 100.00% (*M* = 75.37, *SD* = 21.67), corresponding to ITRs of 9.51 - 128.2 bpm (*M* = 80.41, *SD* = 8.51; **Figure 3*a***). Reliable free communication was deemed too difficult when classification accuracy was less than 80%. Therefore, only participants with high classification accuracy were considered for free communication (*N* = 9/17; *M* = 92.29, *SD* = 2.20). One high accuracy (82.14%) participant elected to undergo retraining rather than free communication, citing frustration with misclassification. The remaining participants underwent retraining (accuracy < 80%; *N* = 8/17; *M* = 57.47, *SD* = 18.13; **Figure 3*a***).

**Figure 2.**
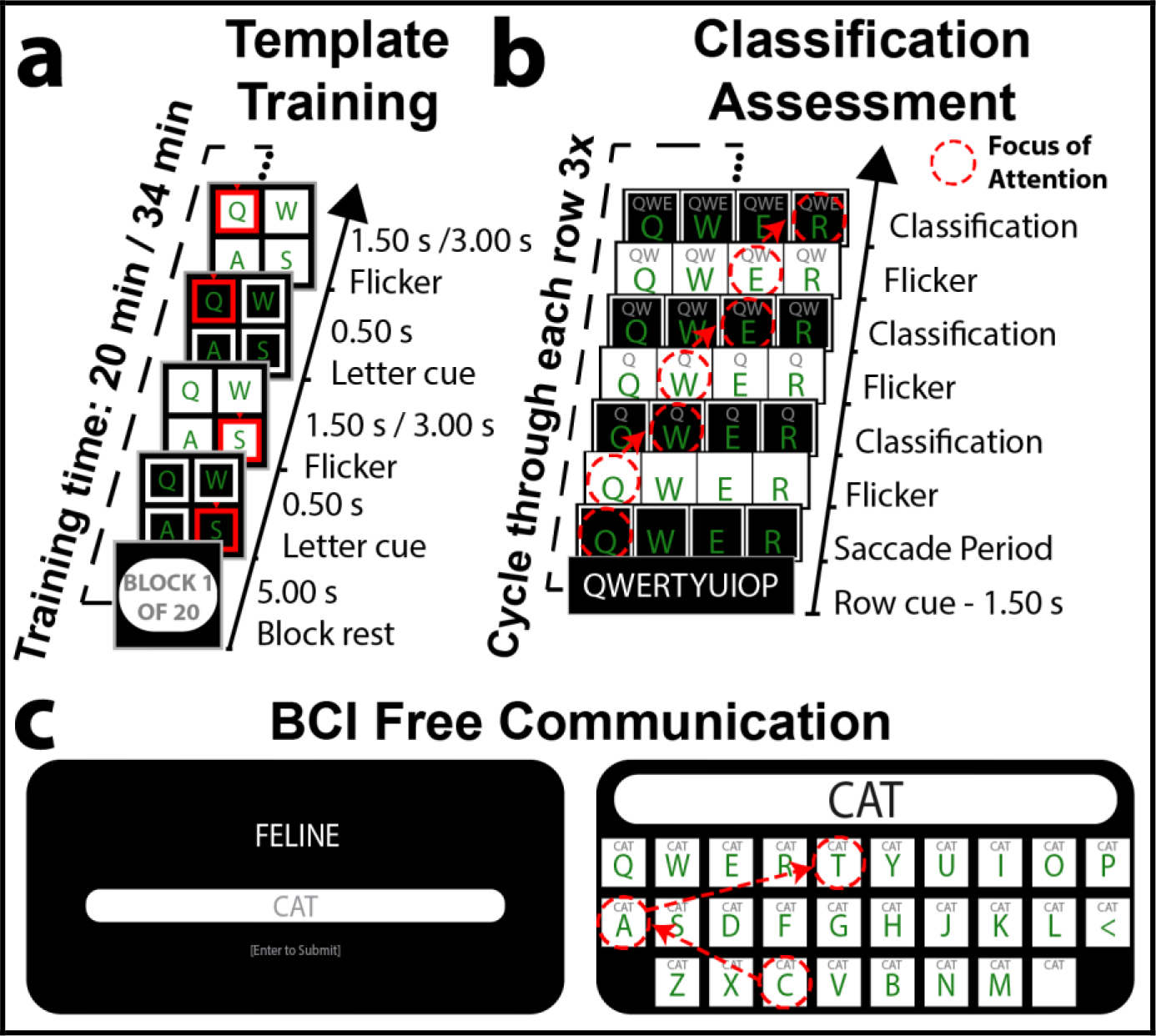
The three phases of Experiment 1, which allowed BCI free communication through prompted free association. (***a***) Template training. Participants (*N* = 17) were cued to focus their attention and gaze on each flickering key in a random order (*N* = 20 repetitions/key). Keys were cued prior to and during flicker. Participants with low classification accuracy (< 80%), determined via QWERTY classification, underwent retraining with an increased flicker duration (3.0 vs. 1.5 s) to improve single-trial SSVEP SNR. (***b***) QWERTY classification. Participants freely BCI typed the complete QWERTY sequence, without guiding cues other than feedback displayed at the top of each key. Flicker-free periods allowed participants 0.75 s to redirect their attention to the next uncued key. The three previous classified letters were displayed at the top of each key, allowing participants to monitor classification while performing keystrokes. (***c***) BCI free communication undertaken by participants with high classification accuracy (> 80%). Prompt words allowed participants to freely associate words/phrases. To assess accuracy, participants entered intended character strings using a manual keyboard before BCI typing. A new prompt was presented when participants either matched the intended character string using BCI, or had entered three times more characters than in the intended string.

The retraining group (*N* = 9/17) completed template generation based on a double flicker epoch to increase SSVEP SNR. The flicker signal was presented twice in sequence 3.0 s), and the ERP of the two epochs (1.5 s each) was treated as the single-trial EEG. Again, performance was evaluated by having participants BCI type the complete QWERTY sequence three times, without guiding cues. A paired samples *t*-test demonstrated that classification accuracy (%) was significantly higher for the double (*M* = 78.04, *SD* = 19.96) than single flicker epoch (*M* = 60.22, *SD* = 18.85; *t*_8_ = -4.34, *p* = .002; **Figure3*b***), a mean improvement of ~18%. This increase in classification accuracy resulted in ITRs (bpm) that did not differ significantly for the double (*M* = 50.41, *SD* = 6.57) and single flicker epochs (*M* = 54.12, *SD* = 8.41; *t*_8_ = 0.75, *p* = .476), despite the much longer flicker duration (× 2). The double flicker epoch resulted in 6/9 participants being classified with greater than 80% accuracy, increasing the system’s suitability for free communication. The classification improvement using double epoch ERPs indicated that filter-bank CCA’s algorithms depend on high SNR and single-trial SSVEP phase consistency.

**Figure 3.**
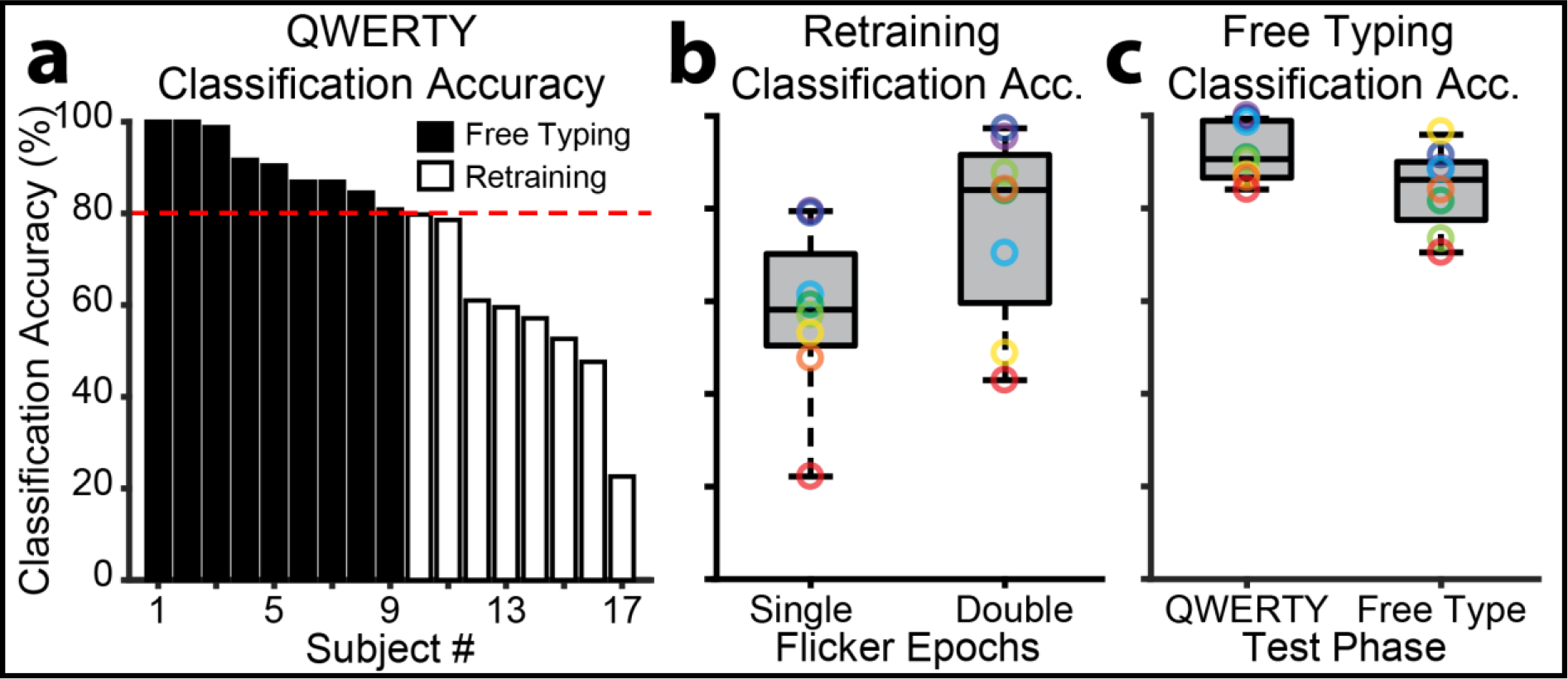
Accuracy during QWERTY classification and free communication. (***a***) Individual differences in QWERTY classification (single flicker epochs: 1.5 s). Participants classified with greater than 80% accuracy (*N* = 8/17) advanced to free communication, with the remaining participants undergoing retraining with a double flicker epoch (3 s). (***b***) Box plot of QWERTY classification for the low accuracy group with 1.5 s single and 3 s double flicker epochs. (***c***) Box plot of performance for the high accuracy group on QWERTY classification and free communication via prompted association. In these and subsequent box plots, coloured rings represent individual participants, the central bars represent the median, outer bars represent the 25^th^ and 75^th^ percentiles, and the whiskers extend to the absolute maxima.

### BCI Free Communication

The primary purpose of Experiment 1 was to determine whether the BCI system was suitable for free communication in naïve users. Therefore, high accuracy participants (*N* = 8/17) freely generated words/phrases that were semantic associates of word prompts (**Figure 2*c***; **Appendix 1**). Participants first manually typed their intended words/phrases using a standard keyboard, and then attempted to replicate these character strings using BCI typing. This allowed us to quantify classification accuracy during free communication.

Participants could successfully freely communicate using the BCI system, generating a large variety of unique words and phrases in response to the prompts (**Appendix 2**). Example prompt/associate pairs include: “GREAT”/“DIM SUM”, “IMPORTANT”/“MUM” and “PLACE”/“IT PUTS THE LOTION ON ITS SKIN”. Participants generated 1-10 words/prompt (*M* = 1.80, *SD* = 0.53), selecting each key at least once ([X] & [Z]) and up to 201 times ([backspace]; *M* = 21.26 selections, *SD* = 8.77). Participants generated words of average complexity (word length: *M* = 5.13 characters, *SD* = 0.22), equivalent to the average length of English words (5.1 characters) ^[47]^. Free communication was perhaps most strongly evident in the ability of many prompts (selected at random) to produce different successfully BCI typed associates across participants (e.g., prompt: “END”; associates: “OF THE DAY”, “FINAL”, “HOLIDAY”, “START”). This indicated that communication depended on the participants’ individual real-time thoughts and that the association task successfully tapped free communication.

Classification accuracy and ITR are often determined by instructing users to repeat phrases or cycle systematically through the keyboard. We included such a classification assessment, with the aim of contrasting free communication performance. A paired samples *t*-test revealed that classification accuracy (%) was significantly lower during free communication (*M* = 84.22, *SD* = 3.09) than on the QWERTY assessment (*M* = 92.29, *SD* = 2.20; *t*7 = 2.96, *p* = .021; see **Figure 3*c***). The performance cost of free communication (~8%) might be partly attributable to increased demands on memory and search during free communication, and indicates that instructed assessments overestimate free communication ITRs.

### Factors Affecting the Feasibility of BCI Free Communication

As classification accuracy varied substantially across participants, we investigated a number of contributing factors. To evaluate SNR, we examined FFT ERP amplitude spectra for each frequency during the (single epoch) template generation phase, undertaken by both the low and high accuracy groups. ERPs for each frequency (*N* = 20 × 1.5 s epochs) were zero-padded to 5.0 s to allow 0.2 Hz spectral resolution and separation of adjacent flicker frequencies (**Figure 4*a***). To evaluate the effect of SNR on classification accuracy, we calculated the difference in FFT amplitude spectra between the high and low accuracy groups (**Figure 4*b***). SSVEP amplitudes were larger for the high compared with low accuracy group, especially for the first three harmonics. The grand mean SSVEP amplitude topographies of the first harmonics revealed maximal amplitudes at occipitoparietal sites, consistent with previous frequency tagging studies of attention ^[48–50]^, with larger amplitudes for the high compared with low accuracy group (**Figure 4*c***). This effect was confirmed statistically: classification accuracy (%) was strongly positively correlated at the between-subjects level with the mean SNR of the first harmonic flicker frequencies (*r*_15_ = 0.84, *p* < .001; **Figure 4*d***). These results demonstrate that SSVEP SNR is critical for reliable free communication.

**Figure 4.**
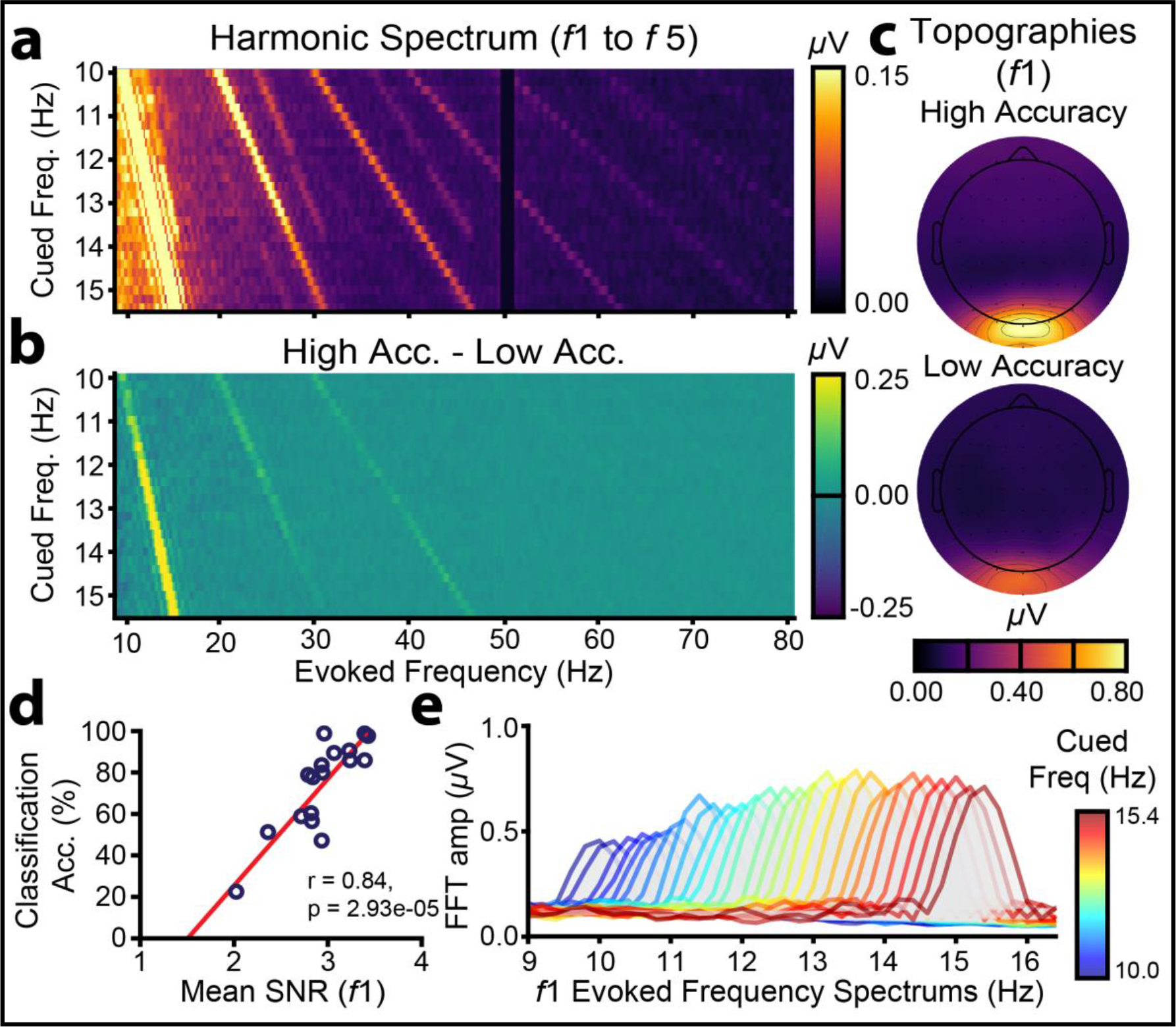
SSVEP SNR. (***a***) Grand mean FFT ERP amplitude spectra across all participants plotted for harmonics 1 - 5. Warmer colours indicate higher SSVEP amplitudes. The colour map is scaled to highlight later harmonics; the slice through the first harmonic in fact extends to 2.10 *μ*V (***b***) Differences in grand mean spectra for the high and low accuracy groups. (***c***) Grand mean SSVEP amplitude topographies at the first harmonic, averaged across all flicker frequencies, plotted separately for the high and low classification accuracy groups. (***d***) Scatter plot of the positive relationship between mean SSVEP SNR at the first harmonics and QWERTY classification accuracy. (***e***) Grand mean FFT amplitude spectra at the first harmonic for each cued frequency during template training.

Related to SSVEP SNR, the range of stimulation frequencies is a free parameter likely to affect classification accuracy. Assessment of the first harmonic SSVEPs revealed that SNR was lower for frequencies in the alpha range (10.0 - 12.0 Hz) than for higher frequencies (12.0 - 15.4 Hz; **Figure 4*e***). This may owe to interference at lower frequencies from phase-misaligned endogenous alpha oscillations ^[51]^. Consistent with this interpretation, classification features 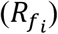 on the QWERTY assessment were higher for *unselected* template frequencies in the alpha range. This effect was apparent only for the low accuracy group (**Figure 5*a***), increasing the likelihood of misclassification for these individuals. Thus, avoiding stimulation frequencies in the alpha range might increase the feasibility of free communication.

Our filter-bank CCA algorithms used multiple stimulation frequency harmonics. *Prima facie*, the inclusion of additional harmonics might improve accuracy, as classification is based on more information. To examine this, we classified the baseline template data ten times, incrementally including an additional harmonic at each step (i.e., [1]: *f*_1_; [2]: *f*_1_, *f*_2_; [3]: *f*_1_, *f*_2_, *f*_3_ …). As depicted in **Figure 5*c***, classification accuracy in fact benefited from fewer rather than more harmonics. To assess this statistically, classification accuracy (%) was submitted to a two-way mixed ANOVA with group (high accuracy, low accuracy) and number of harmonics (1, 2, 3, 4, 5, 6, 7, 8, 9, 10) as factors. There were significant main effects of group (*F*_1,15_ = 25.95, *p* < .001, *η_p_*^2^ = .63) and number of harmonics (*F*_9,21_ = 8.05, *p* = .005, *η_p_*^2^ = .35; Greenhouse-Geisser adjusted, Mauchly’s *W* < .001, *X*^2^ = 263.23, *p* < .001), but no significant group × number of harmonics interaction (*F*_9,21_ = 0.39, *p* = .388, *η_p_*^2^ = .06). Thus, the effect of the number of harmonics was statistically similar for the high and low accuracy groups. Importantly, the main effect of harmonic was better explained by a quadratic (*F*_1,15_ = 37.60, *p* < .001, *η_p_*^2^ = .638) than linear trend (*F*_1,15_ = 4.08, *p* = .062, *η_p_*^2^ = .214), confirming an overall benefit for fewer harmonics. Examination of the individual-level data indicated that classification accuracy peaked for some participants after only one harmonic, while others benefited from the inclusion of up to seven harmonics (see **Table 1** for the optimal number of harmonics for each participant). Classification accuracy was significantly higher when using each individual’s optimal number of harmonics (*M* = 79.55, *SD* = 20.83) than the five harmonics used in real-time (*M* = 75.35, *SD* = 21.69; *t*_16_ = -3.36, *p* = .004; **Figure 5*b***), although modestly so (~4%). Thus, free communication can be facilitated by selecting the optimal number of harmonics separately for each individual.

**Figure 5.**
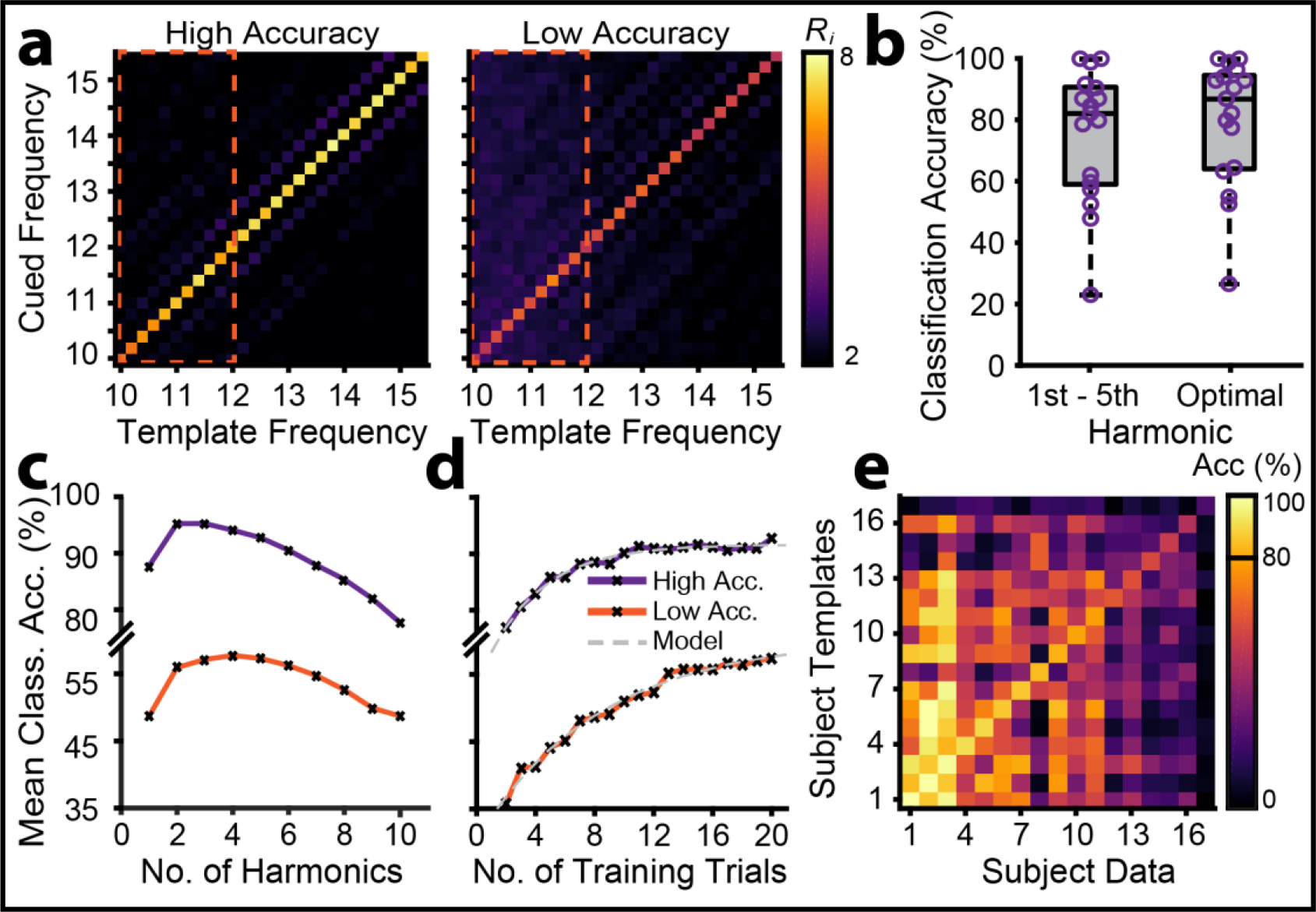
Factors affecting the feasibility of BCI free communication, assessed using QWERTY classification data. (***a***) Grand mean classification features (*R*_*i*_) for each cued frequency for the high and low accuracy groups. For each flicker period, the filter-bank CCA produced a classification feature (*R*_*i*_) representing the correlation between the single-trial EEG and templates at each frequency. Correct classification occurred when *R*_*i*_ was maximal for the template matching the cued frequency. Template frequencies in the alpha range are indicated by the orange bounding boxes (dashed lines). (***b***) Box plots showing the improvement in classification accuracy using the optimal number of harmonics rather than the fixed first five harmonics that were used in real-time. (***c***) Simulated classification accuracy by number of harmonics, plotted for the high and low accuracy groups. (***d***) Simulated classification accuracy by number of training trials, plotted for the high and low accuracy groups. The fitted model is an inverse exponential function. (***e***) Cross-participant classification. To assess template generalisability, each participant’s single-trial EEG was classified using all other participants’ templates. The leading diagonal represents accuracy when participants were classified using their own templates. The leftmost column (read bottom to top) represents the highest accuracy data classified with progressively lower accuracy templates. Similarly, the bottom row (read left to right) represents the highest accuracy templates used to classify progressively lower accuracy data.

**Table 1.**
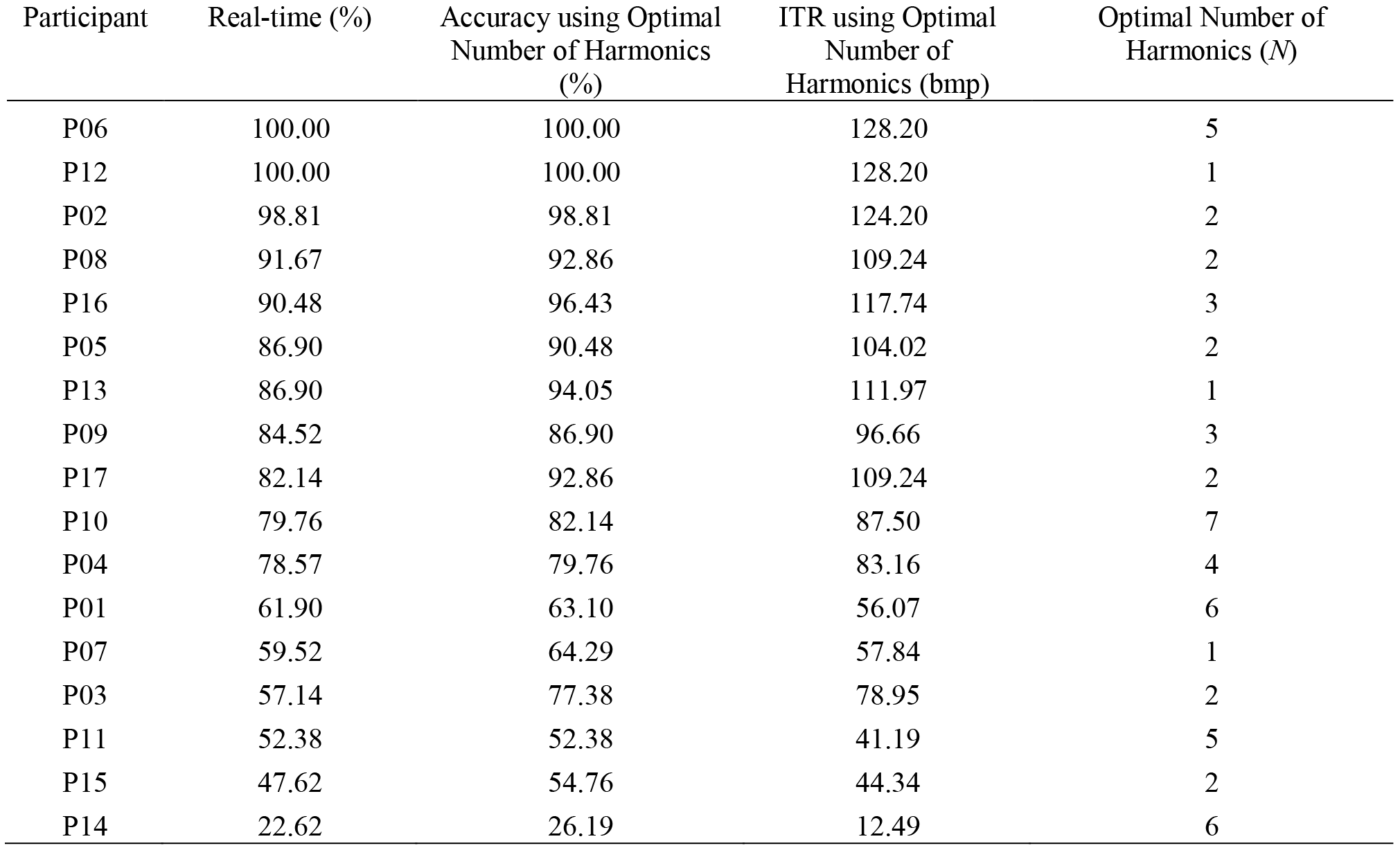
Real-time and offline optimized classification accuracy (%) on the QWERTY classification assessment.

Joint frequency/phase modulated flicker with filter-bank CCA allows high ITRs but can require lengthy individualized calibration. In Experiment 1, template generation required ~18.67 minutes of visual stimulation (2 s/trial × 28 frequencies × 20 trials/frequency). To determine whether shorter calibration is possible, we retrained the filter-bank CCA 20 times for each participant, incrementally including an additional trial for each classification. Using MATLAB’s *fittype* and *fit* functions, which apply the method of least squares, we fitted an inverse exponential function to the resulting classification accuracy curve (**Figure 5*d***):

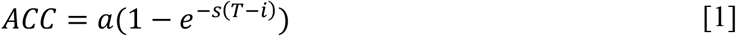

Where *ACC* is classification accuracy, *T* is the number of training trials, *a* is the asymptote, *s* is the scaling factor and *i* is the *x*-axis intercept ^[52]^. On average, the high accuracy group required 12 training trials to reach 99% of their asymptote (90.34% accuracy). In contrast the low accuracy group was projected to require 33 training trials to reach 99% of their asymptote (60.21% accuracy). This suggests that high SNR users require fewer template generation trials for reliable free communication, and that template generation duration can be decreased for these individuals.

While reducing training is desirable, it would be ideal to eliminate training entirely. We therefore assessed whether participants could be cross-classified with other participants’ templates. If cross-classification is feasible, the BCI communication system could be used by new users without individualized calibration. As is evident from the results of the cross-classification analyses (**Figure 5*e***), cross-classification depends on the to-be-classified individual’s own accuracy. Specifically, participants with high classification accuracy were accurately classified with other participants’ templates, while participants with low classification accuracy could not be classified well, even with high accuracy templates. For example, close inspection of **Figure 5*e*** reveals that participant #3 could be classified at over 80% accuracy using 13/17 participants’ templates. There were three cases in which classification accuracy was as good or better using another person’s templates compared with one’s own. This suggests that high SSVEP SNR participants could potentially forego template generation entirely and instead freely communicate from the outset using generic templates.

## Experiment 2: Results

BCI communication systems should be useful not only for expressing thoughts, but also for exchanging thoughts explicitly in conversation with others. To fully evaluate the system’s efficacy for free communication, we therefore extended the system to include an asynchronous two-user messaging interface (**Figure 6**; see also **Video 1** for an early prototype of this interface). Using the default parameters of Experiment 1, leave-one-out classification accuracy on the template training data was 96% (ITR = 131.25 bpm) for participant one (P1) and 98% (ITR = 137.12 bpm) for participant two (P2), suggesting that the classification algorithms were ready for testing under brain-to-brain free communication (~5.7 words/minute). The two participants freely conversed using the BCI messaging interface for ~55 minutes, using an EMG enter key [↲] activated by jaw clenching to complete their messages, view the messaging display and recommence BCI typing. The participants’ conversation focused on recent and upcoming social engagements and their immediate experience using the social interface (see **Appendix 3** for an unedited transcript). In total the two participants generated 68 messages (P1: 33; P2: 35), including 349 words (P1: 170, P2: 179), 1,731 characters (P1: 824, P2: 907) and 283 spaces (P1: 139, P2: 144). Mean word length was 4.2 characters (*SD* = 2.2, range = 1-13), and was similar for the two participants (P1: *M* = 4.0, *SD* = 2.1, range = 1-13; P2: *M* = 4.3, *SD* = 2.3, range = 1-13). To evaluate the conversation’s trajectory during recording, the topic of each message was qualitatively scored (**Figure 7*a***). Nine conversation topics were identified, with each participant contributing at least one message to each topic. Interestingly, the topics overlapped in time: at least two and as many as five topics were ongoing concurrently. This appeared to reflect an emergent feature of the interface, allowing each participant to BCI type continuously, rather than iteratively awaiting their partner’s reply.

**Figure 6.**
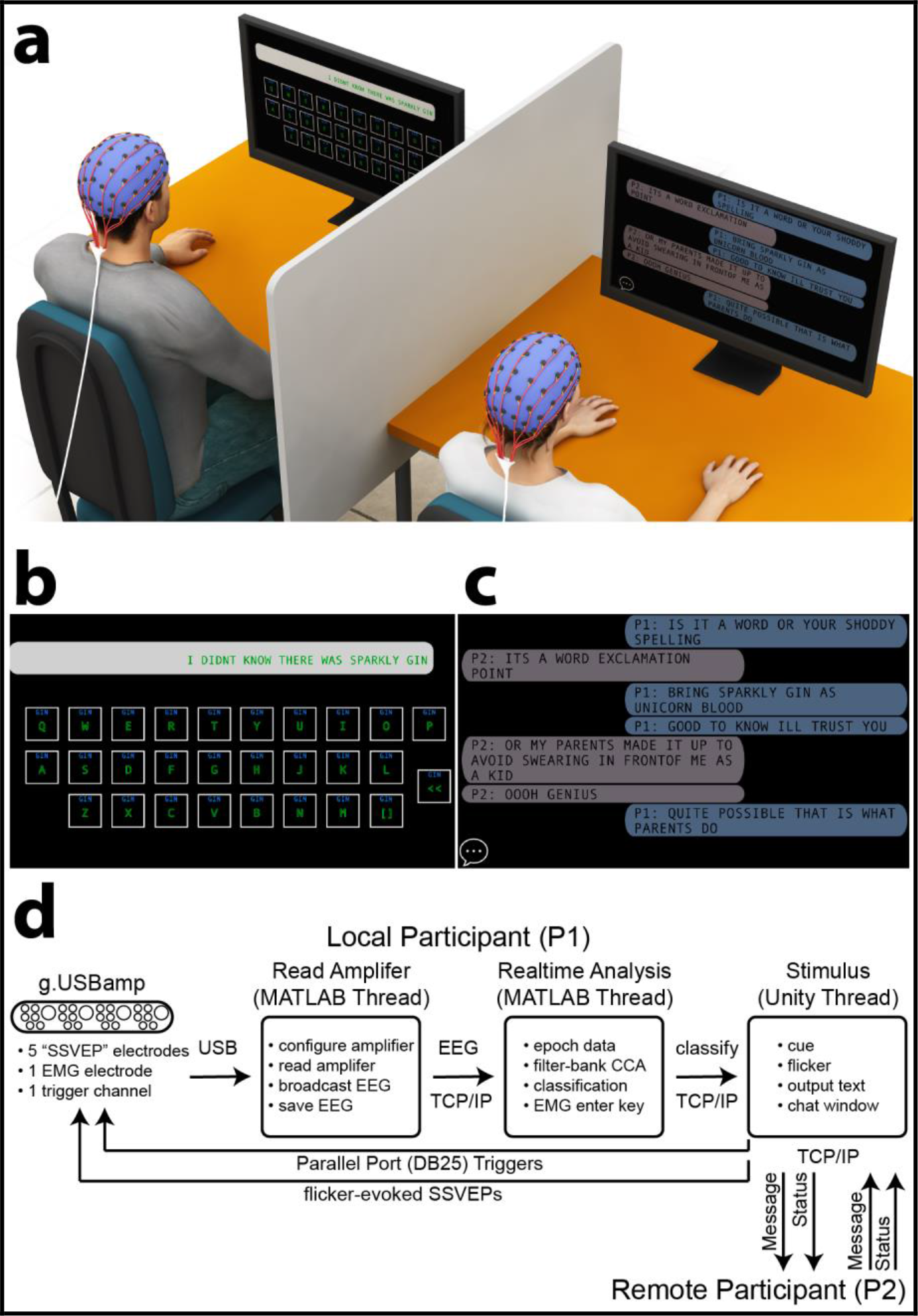
Experiment 2: BCI communication system for brain-to-brain free communication. (***a***) Two experienced participants used an asynchronous messaging interface to have an unprompted free conversation using a system with only six electrodes. (***b***) The keyboard layout was similar to Experiment 1. The interface additionally included an electromyography (EMG) enter key [↲], controlled by detecting jaw clench signals at a frontal scalp electrode, allowing the participants to complete their messages, view the messaging display and recommence BCI typing. (***c***) BCI messaging display. The local participant’s (P1’s) messages are indicated with light blue, and the remote participant’s (P2’s) with light grey. The chat icon is enabled (bottom left corner), indicating that P2 is currently BCI typing, rather than viewing messages. (***d***) Three parallel threads devoted to reading the EEG, real-time analysis and stimulus presentation (indicated by rounded boxes). Inter-thread communication was via parallel port and TCP/IP (indicated by arrows). “Message” indicated the BCI typed text, and “status” indicated whether the remote participant was typing or viewing messages. The flow of information between threads was identical for P2.

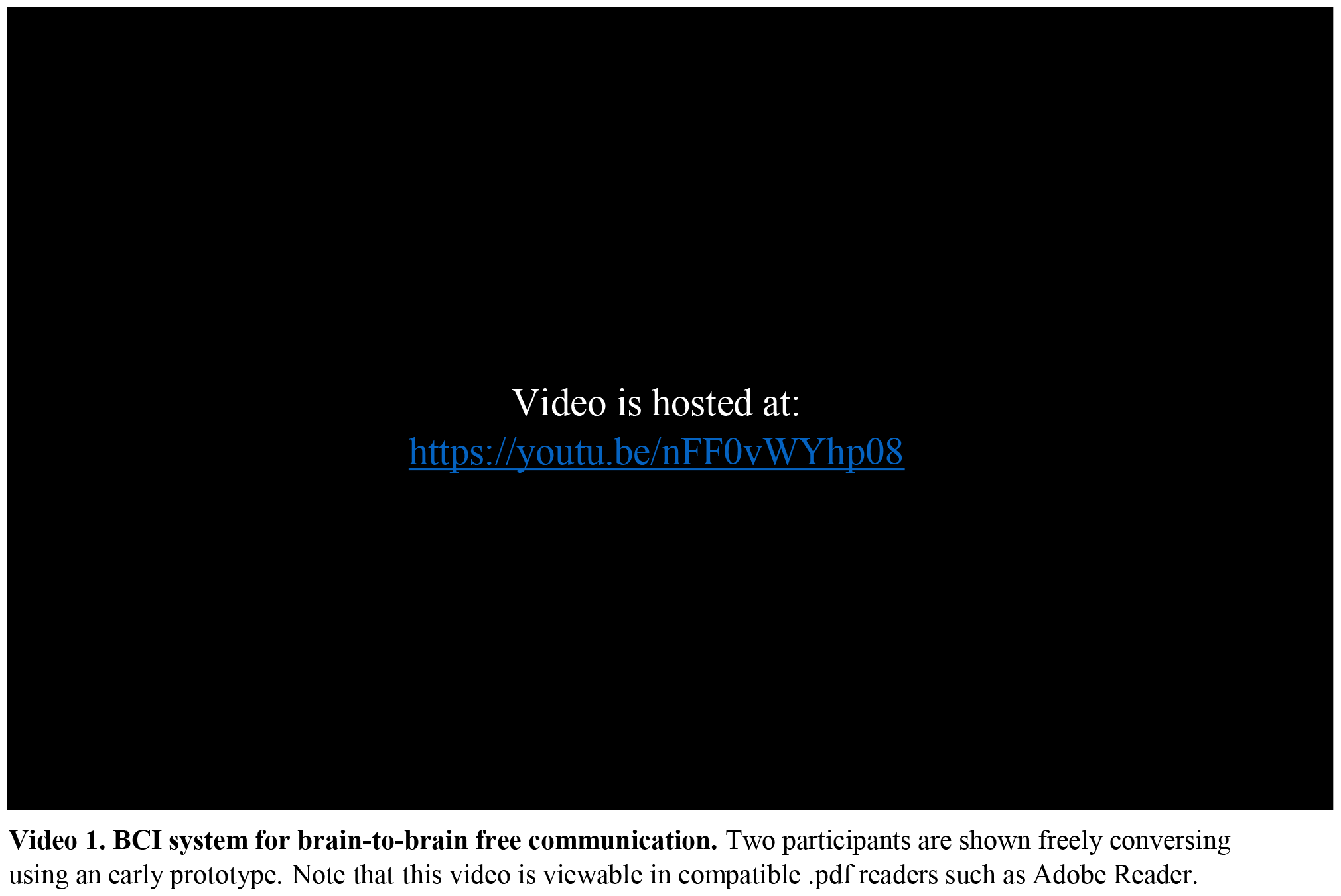

**Figure 7.**
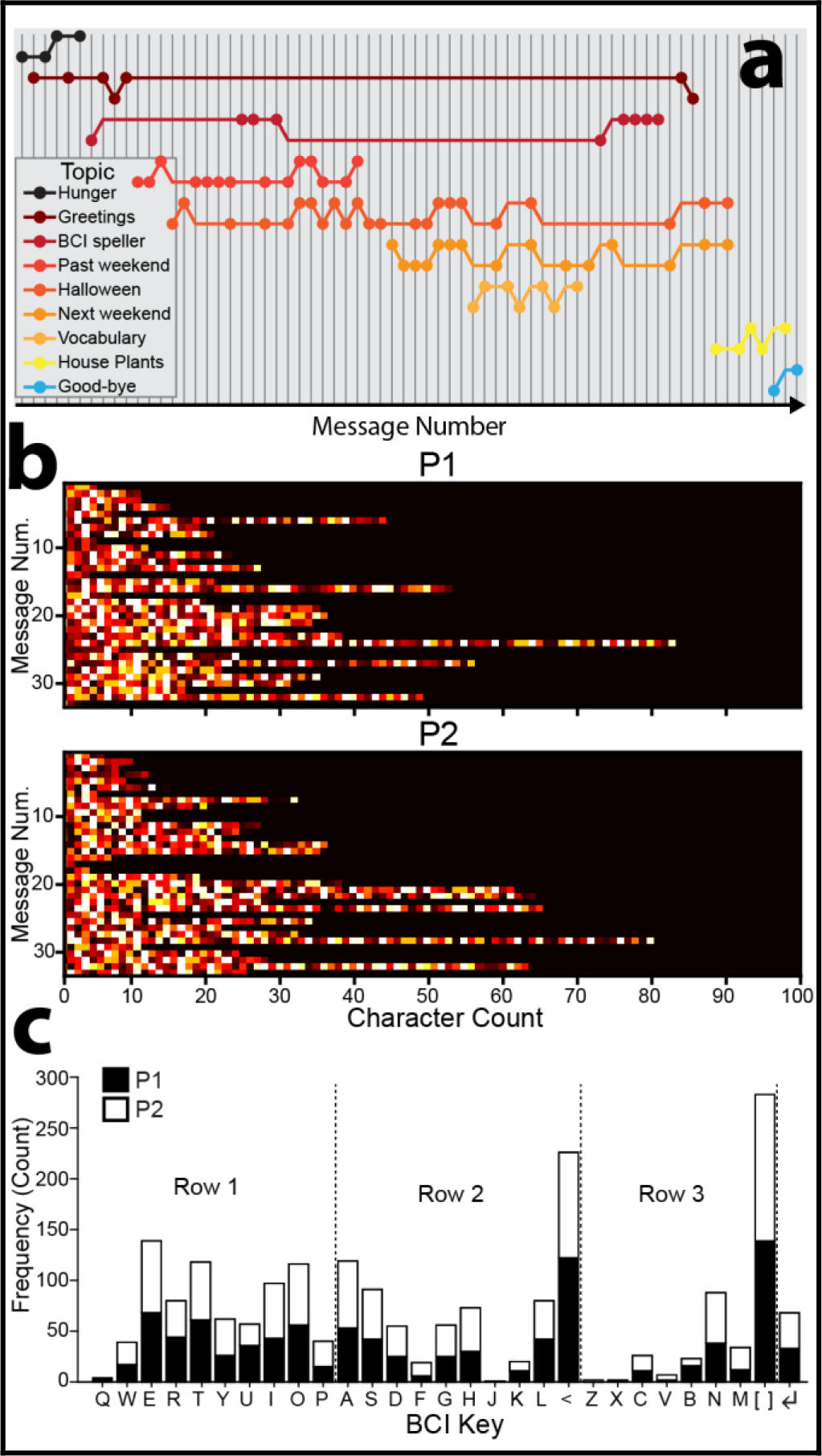
Brain-to-brain free communication results of Experiment 2. (***a***) Streams of brain-to-brain free conversation. Each message was qualitatively scored as belonging to one of nine conversation topics. Each colour represents a topic, and each vertical grey line represents a message. Circles higher in the conversation thread for each topic represent messages sent by P1, while lower circles represent messages sent by P2. (***b***) Messages sent for the two participants (P1 & P2) as a function of ordinal message number and character count. Brighter colours indicate selection of keys later in the keyboard (row three). (***c***) Key selection counts for the two participants, including an EMG enter key [↲] activated by jaw clenching.

For both participants, there was a statistically reliable positive Pearson correlation between message length and ordinal message number (P1: *r*_31_ = .39, *p* = .027; P2: *r*_33_ = .50, *p* = .002; **Figure 7*b***), suggesting that the participants gained confidence in the interface over the course of recording. The participants on average spent 78% of the recording duration BCI typing, with the remaining 22% devoted to viewing typed messages. The considerable proportion of time spent viewing messages indicated that ITRs calculated offline overestimate pure character transfer during free brain-to-brain communication, which naturally entails turn taking.

As depicted in **Figure 7*c***, the backspace and space keys were selected most frequently by both participants, and together the participants used each BCI key at least once. To test whether the two participants differentially selected particular keys, chi-square tests of independence set expected values (*N*) at half the total number of observations for each key. Twenty-four of the 29 keys (including the EMG enter key [↲]) had sufficiently large expected values (*N* > 5) to test statistically, excluding keys [Q], [J], [Z], [X] and [V], which were selected too infrequently. Observed and expected values did not differ significantly for any of the keys (χ^2^s < 3.90, *p*s > .0484; α = .002; Bonferonni α = .05/29 = .002), indicating that key selection was similar for the two participants during free communication. Counting backspace corrections as classification or participant errors, overall classification accuracy during free brain-to-brain communication was 88% (ITR = 98.86 bpm) for P1 and 90% (ITR = 103.01 bpm) for P2, down 8% from training classification. These results show that the BCI messaging interface was suitable for free communication, and that offline classification performance reflects an upper estimate for ITR during free communication.

## General Discussion

Free and open communication is central to modern civilisation, allowing people to convey their thoughts and interface with computers and the internet. While individuals are remarkably adept at operating manual keyboards, a next frontier is communication without manual input. Here, we developed a high performance SSVEP speller based on filter-bank CCA, and examined its feasibility for free communication. In Experiment 1, we tested whether naïve users could maintain rapid typing during prompted free word association. In Experiment 2, we developed a social messaging interface, allowing two experienced users to have an unprompted free conversation. Overall, our results showed that traditional cued typing tests do overestimate free communication ITRs (**Figure 3*c***). However, given individualised interfaces involving sufficient template training trials (**Figure 5*d***) and flicker durations (**Figure 3*b***) and appropriately chosen harmonic (**Figure 5*b-c***) and frequency (**Figures 4*e*** & **5*a***) parameters, the majority of naïve users would be able to freely communicate. The single greatest determinant of free communication success was SSVEP SNR, which showed a strong positive correlation with classification accuracy (**Figure 4*d***). Our results suggest that individuals with high SNR might begin free communication earlier in recording due to the reduction of training trials and the ability to be cross-classified with generic templates (**Figure 5*e***). The successful brain-to-brain free conversation of Experiment 2 (**Appendix 3**) suggested that experienced users with high SSVEP SNR could use the virtual keyboard as naturally as a manual keyboard, albeit more slowly. Message character counts increased for both participants during their free conversation (**Figure 7*b***), suggesting that experienced users can improve their BCI communication efficiency, even when classification parameters remain constant.

Our communication system was made possible by the strong foundations and remarkable recent progress in the field of non-invasive BCI spellers. The first BCI speller, based on the P300 ERP, could reliably classify characters at a rate of 12 bpm (~2.3 characters min^−1^) ^[25]^. Modern spellers using SSVEPs, which offer higher single-trial SNR relative to classical ERPs, can achieve rates of around 146 bpm, an order of magnitude faster than early spellers ^[34; 37–41]^. The seminal development of filter-bank CCA has allowed the report of an unprecedented ITR of 267 bpm, reflecting a forward step in non-invasive neuroimaging ^[21; 38]^. However, these impressive leaps in ITR were generated by typing generic, cued character strings. Ultimately, BCI systems are intended for free communication, which is inherently interactive and spontaneous.

Our studies investigated the useability of BCI systems for free communication. As a first step, we developed three tests with increasing levels of user freedom and expression: (1) QWERTY classification in which users were instructed but not explicitly cued to complete the entire keyboard sequence. In conjunction, we introduced status characters at the top of each key that allowed participants to track classification while entering keystrokes. (2) We developed a prompted free association task in which the ground truth for character intention was established by having users enter the target phrase using a manual keyboard before BCI typing. Free association allowed users to generate their own words and phrases with minimal external input. (3) We introduced free “brain-to-brain” communication allowing users to converse freely, without input from the experimenter. To support free conversation, we developed a social BCI interface that allowed users to view and respond to their conversational partner’s messages, received asynchronously. Our results support the use of joint frequency-phase modulated CCA BCI systems for free communication, but indicate that ultra-high ITRs are not realistic for free communication given current interfaces, which require serial key selection and lack predictive text.

Our results indicate that effective free communication requires a focus on useability rather than fast character selection time. Pilot development indicated that naïve users could not reliably make saccades to the next key with the short trials durations (i.e., 0.5 s flicker/0.5 saccade) employed in previous work ^[21; 38]^, especially during free communication. For these users, short durations were not conducive to free communication due to the overwhelming cognitive load of focusing selective attention on a virtual key, ignoring distraction from adjacent flickering keys, planning successive keystrokes, locating the next key and making backspace corrections for misclassifications. The effect of the added cognitive load during free communication is illustrated by the ~8% reduction in classification accuracy experienced by the eight naïve users of Experiment 1 who progressed from the QWERTY assessment to free communication. Consistent with this result, the two experienced users in Experiment 2 also showed an ~8% reduction in classification accuracy from offline assessment to free communication. Thus, cued and instructed typing tests overestimate free communication ITRs. Reductions in classification accuracy during free communication may have been more drastic if not for our usability improvements, including reducing the number of keys, using a QWERTY layout and displaying classification feedback on each key.

Improving useability also involves improving classification accuracy. Individual differences in classification accuracy were largely attributable to SSVEP SNR. Participants with low SNR were therefore retrained using a double flicker epoch, with classification based on the mean SSVEP of the two epochs. This nearly doubled character selection time, but greatly increased classification accuracy (+18%), allowing ITRs to remain constant. Our results indicate that a focus on useability provides ITRs sufficient and practical for free communication, despite longer character selection times relative to cued spelling. Indeed, as users anecdotally remarked through BCI typing: “I WANT ONE OF THESE ON MY PHONE” and “TYPING WAS NEVER BETTER”. Ultimately, usable interfaces for free communication require serviceable rather than ultra-high ITRs.

A main advantage of communication systems based on filter-bank CCA is that the analysis parameters can be adapted and individualised to optimise performance. For instance, our results showed that the most reliable determinant of classification accuracy was SSVEP SNR. A relatively simple procedure for optimising performance would be to determine SNR across a range of frequencies before template training ^[53]^. This could help select the optimal frequency range, which would be especially advantageous for participants with low SNR in the alpha range. Determining the SSVEP SNR early could allow high SNR participants to proceed immediately to free communication using generic templates, while low SNR participants might undergo double epoch training. Additionally, template training could be optimised by real-time modelling of increases in classification accuracy with additional training trials, which we show is well-characterized by an exponential function that approaches an asymptote. Therefore, real-time evaluation using principled stopping rules might optimise accuracy and minimize training time. Real-time evaluation might also determine the optimal number of harmonics for each individual. Together, these additional measures would optimise performance and reduce training time, improving usability and end-user experience.

The rise of artificial intelligence and virtual/augmented reality has created new opportunities for BCI communication systems beyond their clinical origins. For instance, future systems might allow everyday users to communicate with peers as they navigate virtual worlds, or allow others to discretely compose emails while walking to their next business meeting. Our communication system functions as an early prototype for general-purpose use in naïve users, with many open paths. Development could, as we have, focus on non-invasive sparse electrode systems, which are suited to affordability and portability. Our results demonstrate that well-designed spare-electrode systems can provide high-performance. Real-time classification accuracy was ~89% for brain-to-brain free conversation using only five classification electrodes. Further, individualizing the system can improve classification performance by as much as 18%, as we showed by increasing the stimulus duration, which recommends adaptive interfaces. Adaptive interfaces might maximize efficiency using general-purpose templates, as shown by our cross-classification analysis, or introduce new features such as predictive text ^[54–59]^ or mental imagery decoding^[60]^ to narrow the search space of possible user intentions, increasing efficiency especially for low SNR users.

BCI communication systems have applications hitherto futuristic, made possible by recent advances in signal processing and decoding of neural activity patterns. We have shown that appropriate improvements to existing systems allow major increases in usability, here enabling free communication in naïve users. More specifically, given individually tailored analysis parameters and explicit usability design, filter-bank CCA provides a powerful basis for robust BCI free communication. To explore this possibility further, we recommend that performance appraisals of future systems reflect their intended modes of naturalistic free communication and control.

## METHODS

### Experiment 1: Methods

#### Participants

Seventeen BCI naïve participants (7 males, age *M* = 25.12 years, *SD* = 6.82) volunteered after providing informed written consent and were paid $40. All participants were highly familiar with the QWERTY keyboard layout (typing speed *M* = 231.76 characters min^−1^, *SD* = 52.62 characters min^−1^; ITR = 1152 bpm) but naïve to the speller. The mean typing speed for the group ranks at the 59^th^ percentile of over 46 million runs (https://typing-speed-test.aoeu.eu/). Experiments 1 and 2 were approved by The University of Queensland Human Research Ethics Committee. The participants of Experiment 2 provided consent to have their deidentified data made open access (**Table 2**).

**Table 2.** Online source code and data repositories. Source code includes all template training, real-time classification and visual stimulation routines.

| Description | URL |
| --- | --- |
| Experiment 1/2 Source | https://github.com/davidrosspainter/brain-to-brain-interface/ |
| Experiment 2 Data | https://osf.io/xsq8y/files/ |

#### Overview of BCI Communication System

The system implemented joint frequency/phase modulated flicker paired with a filter-bank CCA (**Figure 1*a-b***). The joint frequency/phase modulated flicker method relies on constant electrophysiological latency across stimulation frequencies, and sets similar flicker frequencies at uncorrelated phases. Filter-bank CCA improves the classification accuracy of standard CCA by using the SSVEP harmonics in combination with the fundamental frequencies. In this study, 28 virtual keys ([A]-[Z], [SPACE] & [BACKSPACE]) were arranged in a QWERTY keyboard layout. Each key was tagged with sinusoidal flicker at a unique frequency/phase (**Figure 1*c***).

#### Experiment Protocol

##### Phase 1: Template training

This procedure was used to generate individualised templates of the neural activity evoked by focusing on each key in the virtual keyboard. A red outline and arrow cued participants to foveate and focus their attention on each key. Each key was cued 20 times in a random order. Keys flickered for 1.5 s followed by a 0.5 s flicker-free period, during which eye movements could be made to the next letter (see **Figure 2*a***). A 5 s rest period followed each cycle through the keys. The recording duration was ~20 minutes, including rests.

##### Phase 2: QWERTY classification

This phase determined classification accuracy using the training templates. Participants focused on each key for one flicker period, cycling through each row from left to right/top to bottom, starting at [Q] and ending at [SPACE]. Participants cycled through the keyboard three times. Importantly, typing was self-directed as no cue was presented to direct participants’ attention to the correct key. Instead, feedback was provided by status characters reflecting the last three classifications, printed at the top of each key (**Figure 2*b***). The flicker-free period was set to 0.75 s. Letter classification took ~0.30 s to compute; thus participants had an additional 0.45 s to redirect their gaze if necessary, which was ample time to complete an eye movement ^[61]^. If classification accuracy in the testing phase was greater than 80%, participants proceeded to free communication (phase 3). Otherwise, participants were deemed unable to reliably communicate and underwent retraining in which two 1.5 s flicker periods were concatenated, with a phase reset after the first 1.5 s. The two 1.5 s epochs were averaged together to form an ERP, which improved the SNR for these individuals.

A challenge for free communication is locating and fixating on the correct key before the onset of the next flicker epoch. Participants were therefore given three minutes to practice using the BCI system before the free communication task began.

##### Phase 3: BCI free communication

This phase assessed the BCI system’s suitability for free communication. Participants were instructed to generate responses to prompt words in a free association task^[45]^. At the beginning of each trial, participants were presented with a prompt word, which was randomly selected from a list of 321 common English words (**Appendix 1;** selected from: https://www.ef-australia.com.au/english-resources/english-vocabulary/top-3000-words/). Participants used a physical QWERTY keyboard to manually type the first word or phrase which came to mind upon seeing the prompt. Participants then attempted to replicate the character string using the BCI system (**Figure 2*c***). When the BCI typed character string matched that submitted using the manual keyboard, the trial ended and a new prompt was presented. If participants were unable to replicate the target character string after entering more than three times the target number of characters, the trial was aborted and a new prompt was presented. Participants performed the free association task for 30 minutes. Classification accuracy was calculated by comparing BCI-entered characters with the target string. Superfluous characters were counted as errors.

#### EEG Recording and Channel Selection

EEG data were sampled at 2048 Hz using a BioSemi Active Two amplifier (BioSemi, Amsterdam, Netherlands) from 64 active Ag/AgCl scalp electrodes arranged according to the international standard 10-20 system for electrode placement in a nylon head cap ^[62]^. The electrode positions were: AF3, AF4, AF7, AF8, AFz, C1, C2, C3, C4, C5, C6, CP1, CP2, CP3, CP4, CP5, CP6, CPz, Cz, F1, F2, F3, F4, F5, F6, F7, F8, FC1, FC2, FC3, FC4, FC5, FC6, FCz, FP1, FP2, FPz, FT7, FT8, Fz, Iz, O1, O2, Oz, P1, P10, P2, P3, P4, P5, P6, P7, P8, P9, PO3, PO4, PO7, PO8, POz, Pz, T7, T8, TP7 and TP8. The common mode sense (CMS) active electrode and driven right leg (DRL) passive electrode served as the ground. EEG data were recorded and streamed to MATLAB using the FieldTrip real-time buffer ^[63]^. EEG data were loaded into MATLAB for template generation using the BioSig toolbox ^[64]^. EEG epochs used in real-time and offline analyses were average referenced, baseline corrected, linearly detrended and notch filtered at 50 Hz.

To determine the optimal EEG channels for template generation and real-time analyses, ERPs at each occipitoparietal electrode site (Iz, O1, Oz, O2, PO7, PO3, POz, PO8, PO4, P1, P3, P5, P7, P9, Pz, P2, P4, P6, P8, P10) were generated for each of the 28 letters/frequencies. These ERPs were zero padded to 5.0 s, allowing 0.2 Hz spectral resolution, and submitted to fast Fourier transforms (FFTs). The four channels showing maximal FFT amplitudes (μV) at each fundamental frequency were retained, such that template generation and real-time classification was based only on the unique channels within this list. These multi-channel data are henceforth referred to as single-trial EEG.

#### Individualized Filter-Bank CCA Classification

The classification procedure determined which frequency/key was most likely selected by finding the template frequency that most strongly correlated with the single-trial EEG, as outlined in **Figure 8**. The analysis evaluated the overall pattern of five weighted correlations between the three input variables: single-trial EEG (*x*; **Figure 8*a***), template sinusoids (*z*; **Figure 8*b***), and template SSVEPs (*y*; **Figure 8*b***), separately for each potential frequency (*f_i_*) and harmonic (*n_j_*):

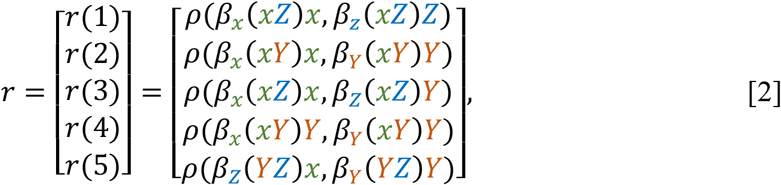

Where *ρ*(*a, b*) represents the weighted correlation between variables *a* and *b*, and *β_c_*(*c, d*) represents the weights of *c* from the canonical correlation of *c* and *d*. The five elements of this vector were combined for each harmonic to form a single selection feature, preserving the sign of each weighted correlation:

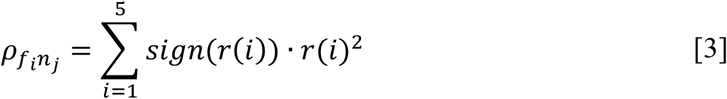

The weighted sum of squares across harmonics was computed as the final feature for target frequency identification:

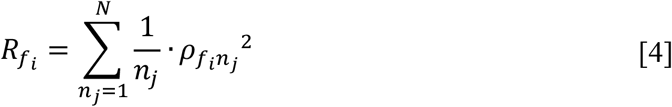

This classification feature 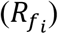 was calculated for each frequency. The frequency/key for which the value was maximal was identified as the selected frequency/key (**Figure 8*c***).

**Figure 8.**
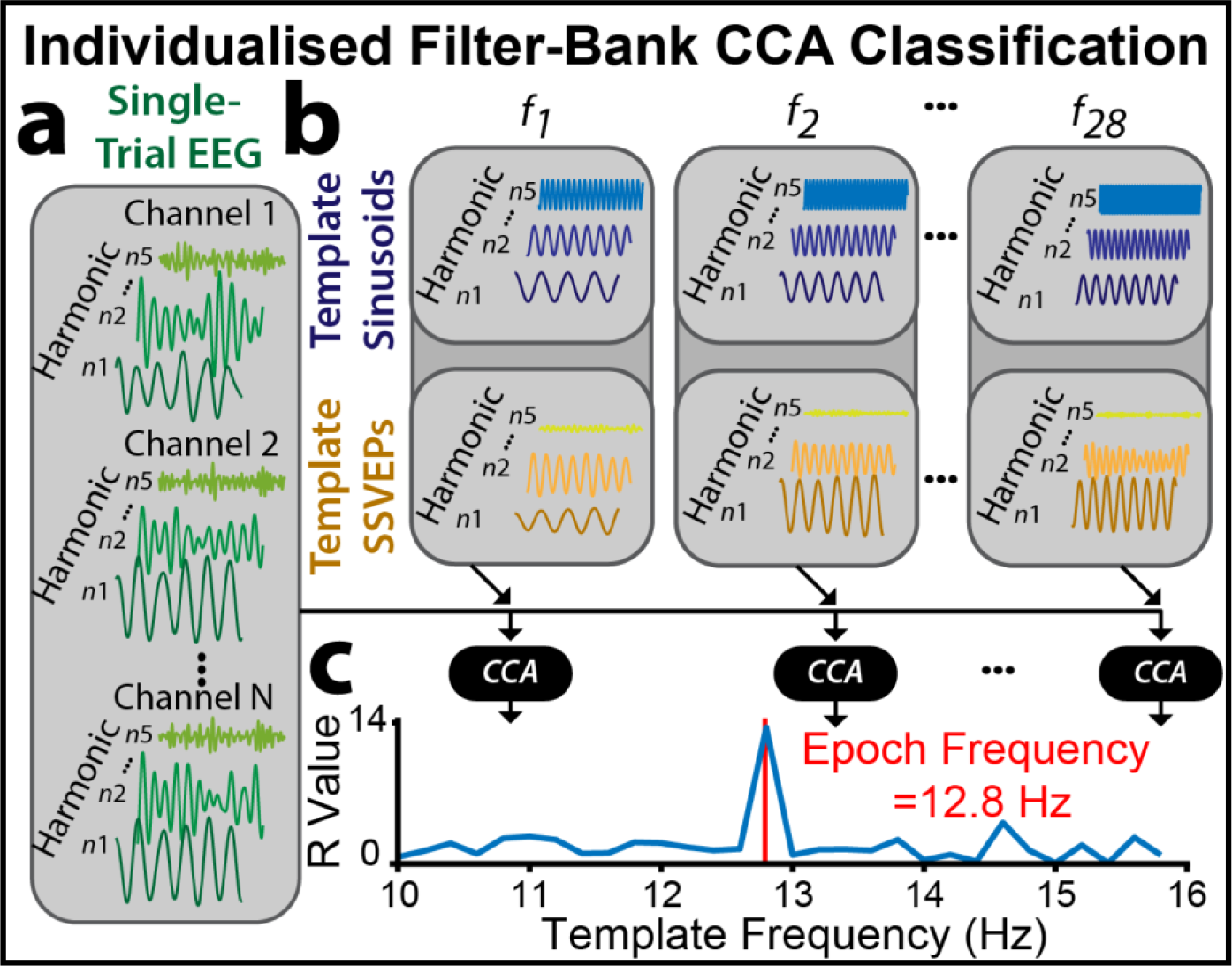
Individualised filter-bank CCA classification. (***a***) Following each flicker period, the corresponding single-trial EEG from the occipitoparietal electrodes with the highest SSVEP training amplitudes were bandpass filtered to the first five harmonic ranges. (***b***) Training resulted in template SSVEPs and sinusoids, reflecting mean signals for each of the 28 flicker frequencies and first five harmonics (see **Methods: Template Training** for a complete description). (***c***) Filter-bank CCA evaluated the overall pattern of five weighted correlations between single-trial EEG, template sinusoids and template SSVEPs. The filter bank CCA produced a final outcome feature for each frequency 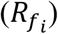, representing the degree of similarity between the single-trial EEG and templates. The frequency with the highest similarity was classified as the selected frequency/key.

#### Template Training

##### Overview

The single-trial EEG recorded during the template training phase was used to create individualised templates of the neural activity evoked by focusing attention on each key. Template signals consisted of 140 (5 harmonics × 28 frequencies) SSVEPs and 140 sinusoids matching in frequency and phase. To increase classification accuracy, the first 0.25 s were excluded, as this reflected a frequency non-specific evoked response.

##### Template bank of SSVEPs

SSVEPs were constructed by averaging the single-trial EEG across the 20 cued trials for each frequency. SSVEPs were bandpass filtered to five harmonic ranges (*n*_1-5_) using a 4^th^ order Butterworth filter:

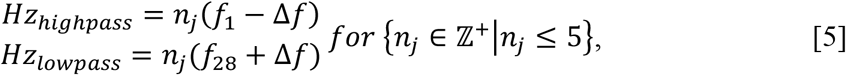

##### Template bank of sinusoids

The procedure for generating template sinusoids matching the SSVEP frequencies/phases is outlined in **Figure 9**. SSVEPs are apparent in the EEG after an onset delay (*τ*; **Figure 9*a***), which potentially varies across individuals and frequencies. The sinusoid phase was therefore calculated separately for each SSVEP frequency and harmonic. An initial bank of potential sinusoids was constructed for each frequency (*f_i_*) at each harmonic range (*n_j_*), including 20 distinct phases (*φ_k_*) from 0 to 1.9 π (**Figure 9*b***):

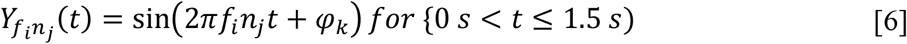

The optimal phase for classification was chosen separately for each frequency and harmonic. To determine the optimal phase, all single-trial epochs corresponding to frequency *f*_*i*_ were bandpass filtered to the harmonic range *n*_*j*_ (**Figure 9*a***). The filtered epochs were correlated (using CCA) with sinusoids at all flicker frequencies and phases (0 - 1.9 π) at harmonic range *n*_*j*_ (**Figure 9*c***). For the CCA, the sinusoid was the univariate measure and the single-trial EEG was the multivariate measure. For each filtered epoch (1 - 20) at frequency *f*_*i*_ and sinusoid phase (0 - 1.9 π), we compared the strength of the canonical correlation (*r*) with each sinusoid frequency (harmonics of 10 - 15.4 Hz). If the maximum correlation for a given epoch was with a sinusoid at the input frequency (*f*_*i*_), the corresponding sinusoid phase was scored as “correct” for that epoch. However, if the maximum correlation was with any other sinusoid frequency, the corresponding phase was scored as “incorrect” (**Figure 9*d***). This allowed us to determine the “accuracy” for each phase by tallying across the 20 epochs (**Figure 9*e***). The phase with the highest accuracy was chosen to be the sinusoid phase for frequency *f*_*i*_ and harmonic *n*_*j*_. In summary, for each frequency and harmonic, we chose the sinusoid phase that maximised classification accuracy by correlating most strongly with SSVEPs at the input frequency during training.

**Figure 9.**
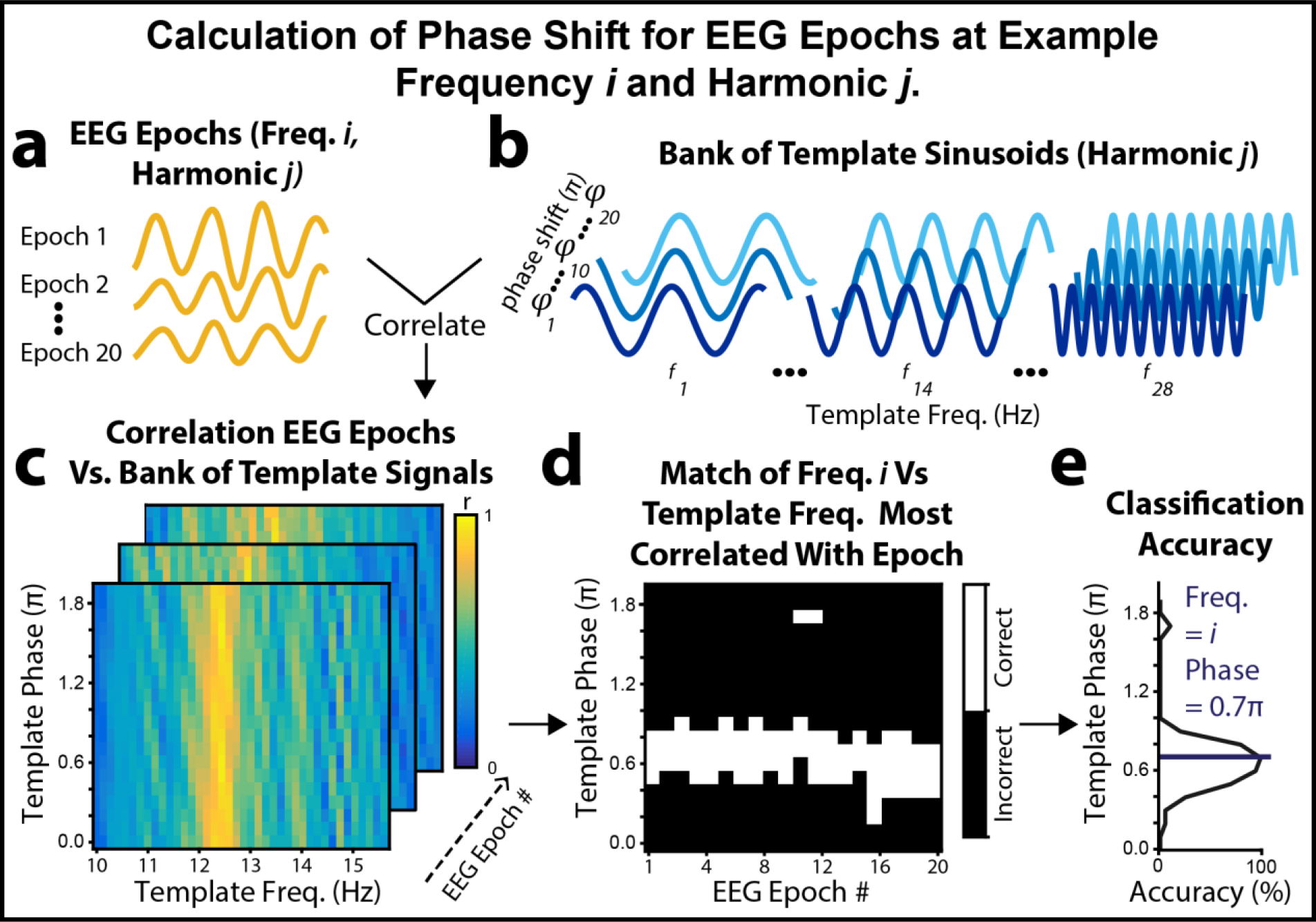
Procedure for generating the template bank of sinusoids with optimal classification phases, calculated separately for each frequency (*i*) and harmonic (*j*; five harmonics in total). (***a***) Single-trial EEG corresponding to cued flicker periods of template training was bandpass filtered to harmonic ranges that encompassed the lowest and highest frequencies for that harmonic (i.e., *n_j_f_1_* - *n_j_f_28_*; e.g., 1*f*: 9.8 - 15.6 Hz). The single trial EEG was canonically correlated with (***b***) each potential template sinusoid (phases 0.0 - 1.9π), resulting in (***c***) canonical *r* values representing the correlation between each single-trial epoch and template sinusoids at each phase. (***d***) For each epoch, the analysis identified the maximally correlated template sinusoid frequency. If this frequency was the cued frequency (*i*), the corresponding sinusoid phase was coded as “correct”, otherwise the phase was coded as “incorrect”. (***e***) The mean classification accuracy across epochs determined the optimal phase of the sinusoid at each frequency and harmonic. In this example, a sinusoid phase of 0.7π maximized classification accuracy.

#### Stimulus Presentation

Each key (*i*) was tagged with sinusoidal flicker at a unique frequency (*f_i_*)/phase (*φ_i_*), with luminance as a function of time (s) since flicker onset (see **Figure 1**):

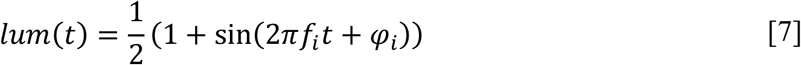

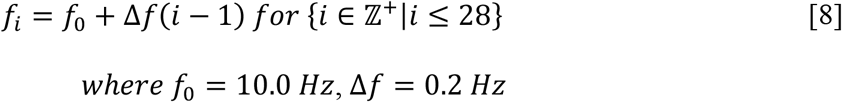

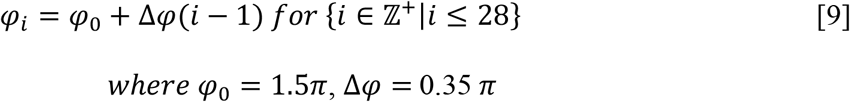

Keys subtended 3.7°^2^ of visual angle, and 1.1° separated adjacent keys. Keys were outlined by a 0.1° white border. Key characters were presented in green 50 pt. Andale Mono font. During training, keys were cued by a red outline and arrow. A white textbox (43.2° × 4.0°) appeared at the top of the display. BCI typed text appeared in this textbox (100 pt. font). During the QWERTY classification assessment and free BCI communication phases, the three last classified characters were printed at the top of each key (40 pt. font), allowing participants to keep track of their BCI typing.

Stimuli were presented at a viewing distance of 57 cm on a 24-inch ASUS VG248QE LCD monitor running at 1920 × 1080 at 144 Hz using the Cogent 2000 Toolbox (http://www.vislab.ucl.ac.uk/cogent.php) running in MATLAB R2016b (64-bit) under Windows 10 (64-bit). The computer contained an Intel Xeon E7-4809 v2 CPU and NVIDIA QUADRO M4000 GPU. The experiment was conducted in a darkened room and participants’ head positions were stabilized with a chin rest. Eye tracking data were also collected for Experiment 1 but are beyond the scope of the present report, which focuses on free communication using BCI alone.

### Experiment 2: Methods

Two experienced BCI participants (ages of 25 and 33 years) were instructed to have a free conversation using their brain activity (**Figure 6a**). Both participants had previously completed three pilot runs in the development of the messaging interface. The two participants viewed separate displays and used BCI systems running on separate computers, with BCI typed messages sent across the local area network via TCP/IP. In addition to the 28 flickering keys of Experiment 1 (**Figure 6b**), we introduced an EMG enter key [↲] based on electrical potentials evoked by jaw clenching. The enter key allowed the two participants to complete their entries and to view the sentences/phrases most recently entered by their conversational partner. A chat icon indicated whether the second participant was currently BCI typing (icon enabled) or viewing messages (icon disabled; **Figure 6c**). The two participants used the enter key at will to resume typing. The EMG analysis involved FFTs performed on the most recent 1 second of data (spectral resolution: 1 Hz) recorded from the frontal scalp electrode FPz. The [↲] key was deemed selected if mean FFT amplitude (*μ*V) in the range of 50-100 Hz exceeded the criterion of 4.0 or 4.5 *μ*V, set separately for the two participants. A cool-down period required that successive enter keystrokes were separated by > 4 s.

The software consisted of three parallel threads devoted to reading from the amplifier, real-time analysis and stimulus presentation (**Figure 6d**). Communication between threads was via parallel port (DB25) triggers and TCP/IP using the FieldTrip real-time buffer ^[63]^. The BCI keyboard and message display were presented using the Unity game engine (Version 2018.2.8f1, Unity Technologies) running at 144 Hz and 1920×1080 pixels with VSync enabled on an NVIDIA GeForce GTX 1080 GPU. The FieldTrip buffer was managed in Unity using a C# API (https://github.com/georgedimitriadis/androidfieldtripbufferinunity). Template generation and real-time classification were performed using eight workers controlled via MATLAB 2017b’s Parallel Computing Toolbox (64-bit) running on an Intel(R) Xeon(R) W-2145 CPU @ 3.70GHz CPU. The calculations underlying template generation required ~2 minutes of processing. Real-time classification occurred during the flicker-free period and took on average ~259 ms (mean of *N* = 1340 classifications; *SD* = 60 ms, range: 135-408 ms).

EEG was sampled at 1200 Hz from g.USBamp amplifiers (one for each participant; g.tec Medical Engineering, GmbH, Austria) from six active gel g.SCARABEO Ag/AgCl scalp electrodes connected to a g.GAMMAbox and arranged in a g.GAMMAcap according to the international standard 10-20 system for electrode placement (Oostenveld and Praamstra, 2001). SSVEPs at five occipitoparietal electrodes (Iz, O1, O2, Oz, POz) were used to create training templates and for real-time classification of the attended BCI key. As noted, electrode FPz was used for the EMG enter key [↲]. The ground electrode was positioned at FCz, and the reference electrode was attached to the left earlobe via a clip. Data were band-pass (1 - 100 Hz) and notch (50 Hz) filtered in real-time at the hardware level. EEG signal quality was established by inspection of the real-time traces visualized by MATLAB’s *dsp.TimeScope* object. The amplifiers were controlled using the g.tec NEEDaccess MATLAB API V1.16.00.

The real-time classification algorithms were identical to those of Experiment 1 (including the use of five harmonics), with the exceptions that the two participants completed 15 (rather than 20) template training blocks (420 trials; 14 minutes) and that SSVEP ERP templates and real-time classification were based on five (rather than four) occipitoparietal electrodes. As in Experiment 1, the flicker period was 1.50 s, and the flicker-free period was 0.50 s for template training and 0.75 s for free communication. The flicker frequencies/phases were identical to those of Experiment 1. The sinusoidal luminance modulation underlying flicker was calculated based on time elapsed from the onset of the first flicker frame using the *System.Diagnostics.Stopwatch* C# class. Frame rates during flicker were confirmed to be stable at 144 Hz using Unity-recorded flip times (*Time.deltaTime*), amplifier-recorded inter-trigger spacing and photodiode measurements. Stimuli were presented on ASUS VG248QE LCD monitors at a viewing distance of 49 cm. The experiment was conducted in a darkened room, and the two participants were separated via a partition.

## Acknowledgements

This research was supported by an Early Career Researcher Seed Funding Grant awarded to DRP by the School of Psychology at The University of Queensland. AIR and DRP were supported by the Australian Research Council (ARC) Special Research Initiative Science of Learning Research Centre (SR120300015). JBM was supported by an ARC Australian Laureate Fellowship (FL110100103) and the ARC Centre of Excellence for Integrative Brain Function (ARC Centre Grant CE140100007).

## Appendix 1. Pool of prompt words for the free association task of Experiment 1

**Appendix 1. Pool of prompt words for the free association task of Experiment 1.**
|  |  |  |  |  |  |  |  |
| --- | --- | --- | --- | --- | --- | --- | --- |
| ABLE | CHILDHOOD | FATHER | HOT | MAYBE | PLAY | SIMPLE | WHITE |
| ACTUALLY | CITY | FEAR | HOUR | MEAN | PLEASURE | SIMPLY | WHOLE |
| AHEAD | CLARITY | FEDERAL | HOUSE | MEDICAL | POINT | SINGLE | WILL |
| AIR | CLEAR | FEEL | HUGE | MEMBER | POLITICAL | SLEEP | WISDOM |
| ALONE | CLEARLY | FINAL | HUMAN | MIGHT | POOR | SMALL | WOMAN |
| ALREADY | CLOSE | FINALLY | IDEA | MILITARY | POPULAR | SOCIAL | WORD |
| AMERICAN | CLOSE | FINANCIAL | IMPORTANT | MINUTE | POSSIBLE | SOMETIMES | WORK |
| APPETITE | COLD | FIND | INDEED | MOMENT | POWER | SOON | WORK |
| AREA | COLDNESS | FINE | INFORMATION | MONEY | PRESIDENT | SORROW | WORLD |
| AROUND | COME | FORCE | INSTEAD | MONTH | PRETTY | SPECIAL | WOULD |
| ART | COMMON | FOREIGN | INTERNATIONAL | MORNING | PRIVATE | START | WRONG |
| ASK | COMMUNITY | FORWARD | ISSUE | MOTHER | PROBABLY | STATE | YEAR |
| AVAILABLE | COMPANY | FREE | JOB | MOVE | PROBLEM | STORY | YOUNG |
| AWARENESS | COULD | FREEDOM | JUSTICE | MOVEMENT | PROGRAM | STRONG |  |
| BACK | COUNTRY | FRIEND | KEEP | MUST | PUBLIC | STUDENT |  |
| BAD | CULTURAL | FRIENDSHIP | KID | NAME | PUT | STUDY |  |
| BEAUTY | CURRENT | FULL | KIND | NATIONAL | QUESTION | STUPIDITY |  |
| BECOME | DARK | GAME | KNOW | NATURAL | QUICKLY | SUDDENLY |  |
| BEFORE | DAY | GENERAL | LATE | NEARLY | QUITE | SURE |  |
| BEGIN | DEAD | GENEROSITY | LAW | NEED | RATHER | SYSTEM |  |
| BELIEF | DEMOCRATIC | GET | LEAVE | NEW | READY | TAKE |  |
| BELIEVE | DIFFERENT | GIRL | LEFT | NICE | REAL | TALK |  |
| BEST | DIFFICULT | GIVE | LEGAL | NIGHT | REASON | TEACHER |  |
| BEST | DIRECTLY | GOOD | LESS | NO | RECENT | TEAM |  |
| BETTER | DOOR | GOODNESS | LET | NUMBER | RECENTLY | TELL |  |
| BETTER | EARLY | GOVERNMENT | LEVEL | OF COURSE | RED | THAT |  |
| BIG | EARLY | GREAT | LIFE | OFFICE | RESEARCH | THING |  |
| BLACK | EASY | GROUP | LIKE | OK | RESULT | THINK |  |
| BODY | ECONOMIC | GUY | LIKELY | OLD | RIGHT | THUS |  |
| BOOK | EDUCATION | HAND | LINE | ONLY | RIGHT | TIME |  |
| BOTH | EITHER | HAPPEN | LITTLE | OPEN | RIGHT | TOGETHER |  |
| BRAVERY | ELSE | HAPPINESS | LIVE | OTHER | ROOM | TONIGHT |  |
| BRING | END | HAPPY | LOCAL | OTHERS | RUN | TRADITIONAL |  |
| BUSINESS | ENOUGH | HARD | LONG | PARENT | SCHOOL | TRU |  |
| CALL | ENTIRE | HARD | LOOK | PART | SEE | TRY |  |
| CALM | ENVIRONMENTAL | HEAD | LOT | PARTICULARLY | SEEM | TURN |  |
| CAN | ESPECIALLY | HEALTH | LOW | PARTY | SERIOUS | USE |  |
| CAR | EVENTUALLY | HEAR | LUCK | PAST | SERVICE | USUALLY |  |
| CASE | EXACTLY | HELP | LUXURY | PEOPLE | SHORT | VARIOUS |  |
| CENTRAL | EYE | HIGH | MAIN | PERHAPS | SHOULD | WANT |  |
| CERTAIN | FACE | HISTORY | MAJOR | PERSON | SHOW | WAR |  |
| CERTAINLY | FACT | HOLD | MAKE | PERSONAL | SIDE | WATER |  |
| CHANGE | FAMILY | HOME | MAN | PHYSICAL | SIGNIFICANT | WAY |  |
| CHILD | FAST | HOME | MAY | PLACE | SIMILAR | WEEK |  |

## Appendix 2. Freely associated and BCI typed words and phrases in Experiment 1

Accuracy indicates whether the BCI typed word/phrase matched (1) or mismatched (0) the text manually entered using a physical keyboard, which provided the ground truth for communication intent.

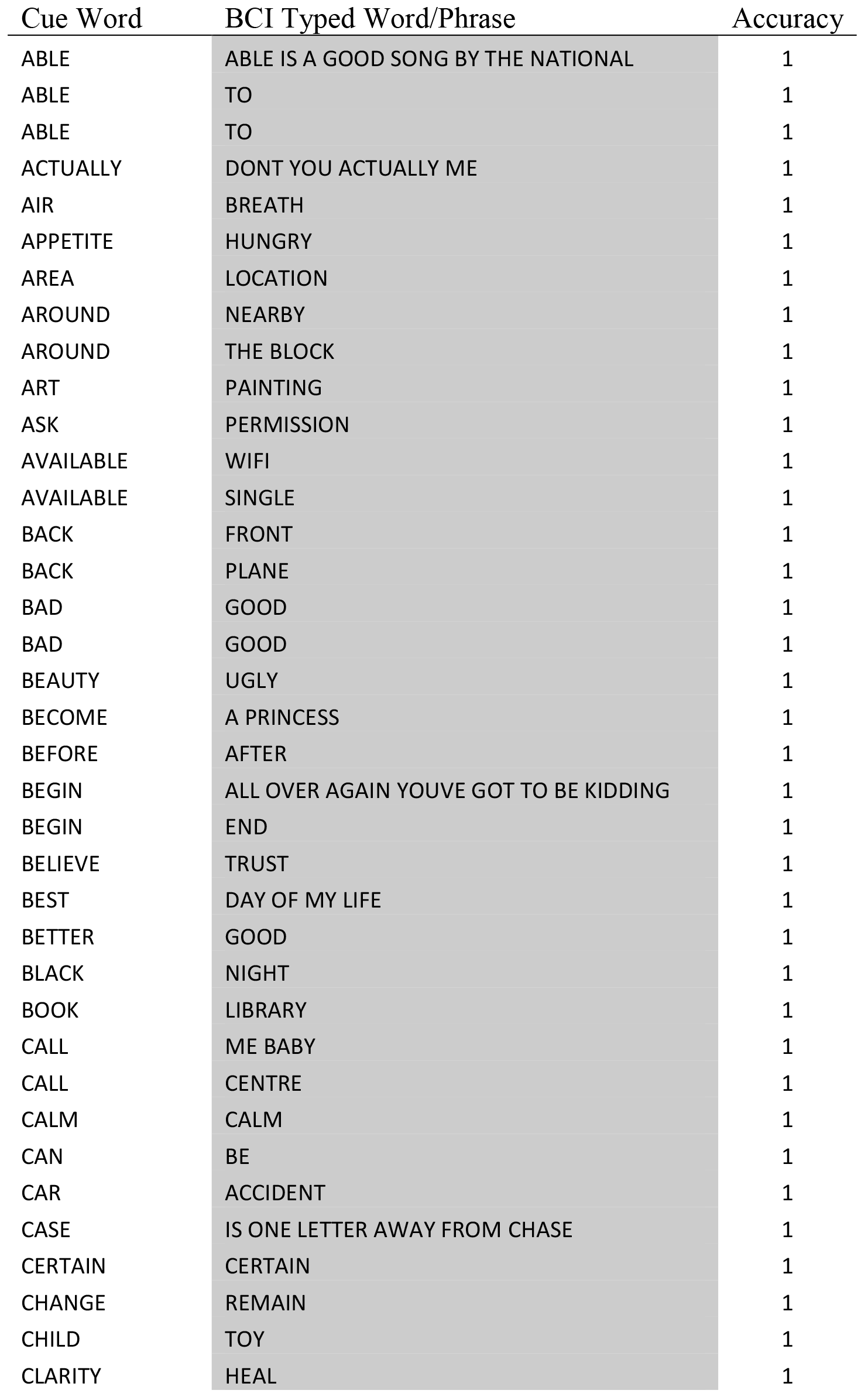

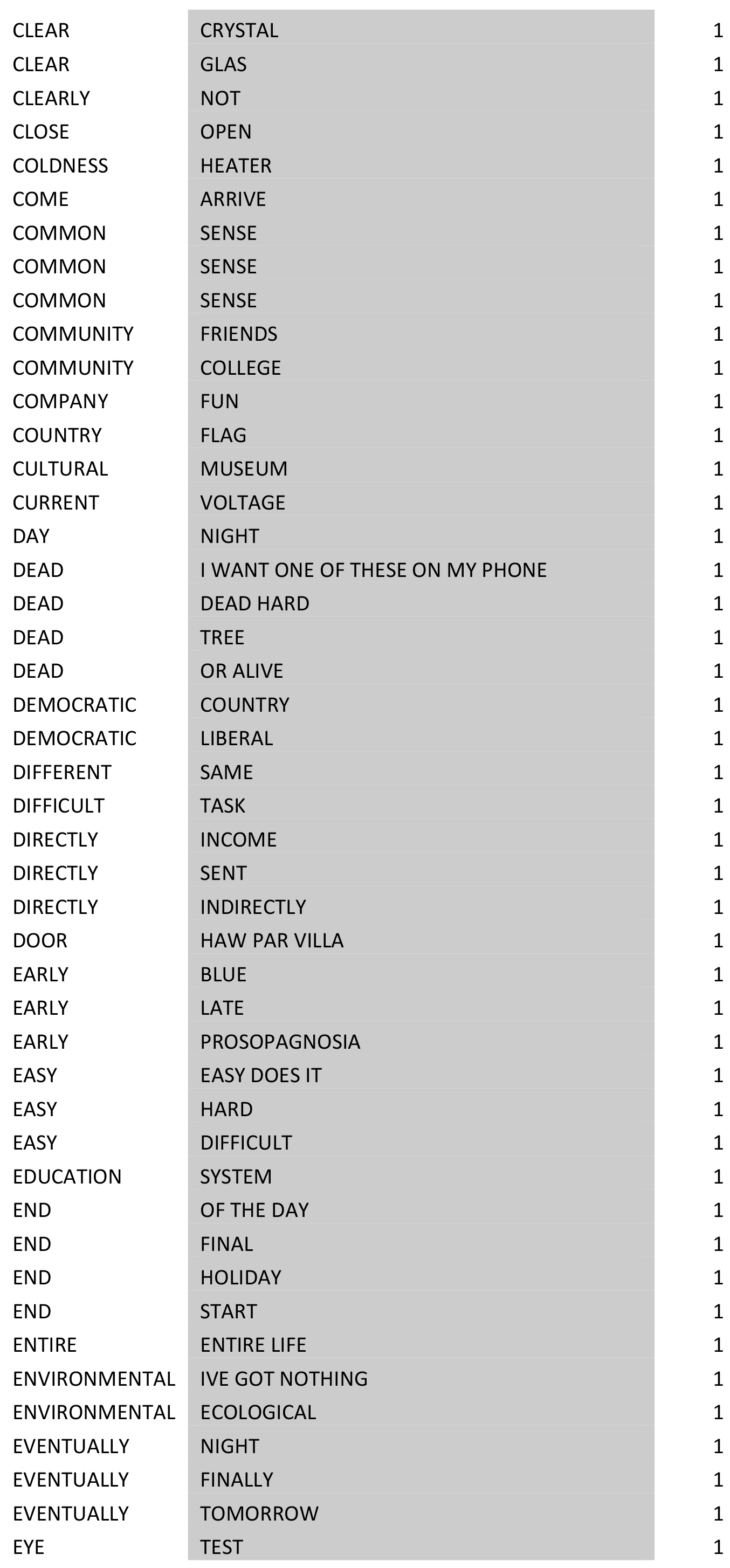

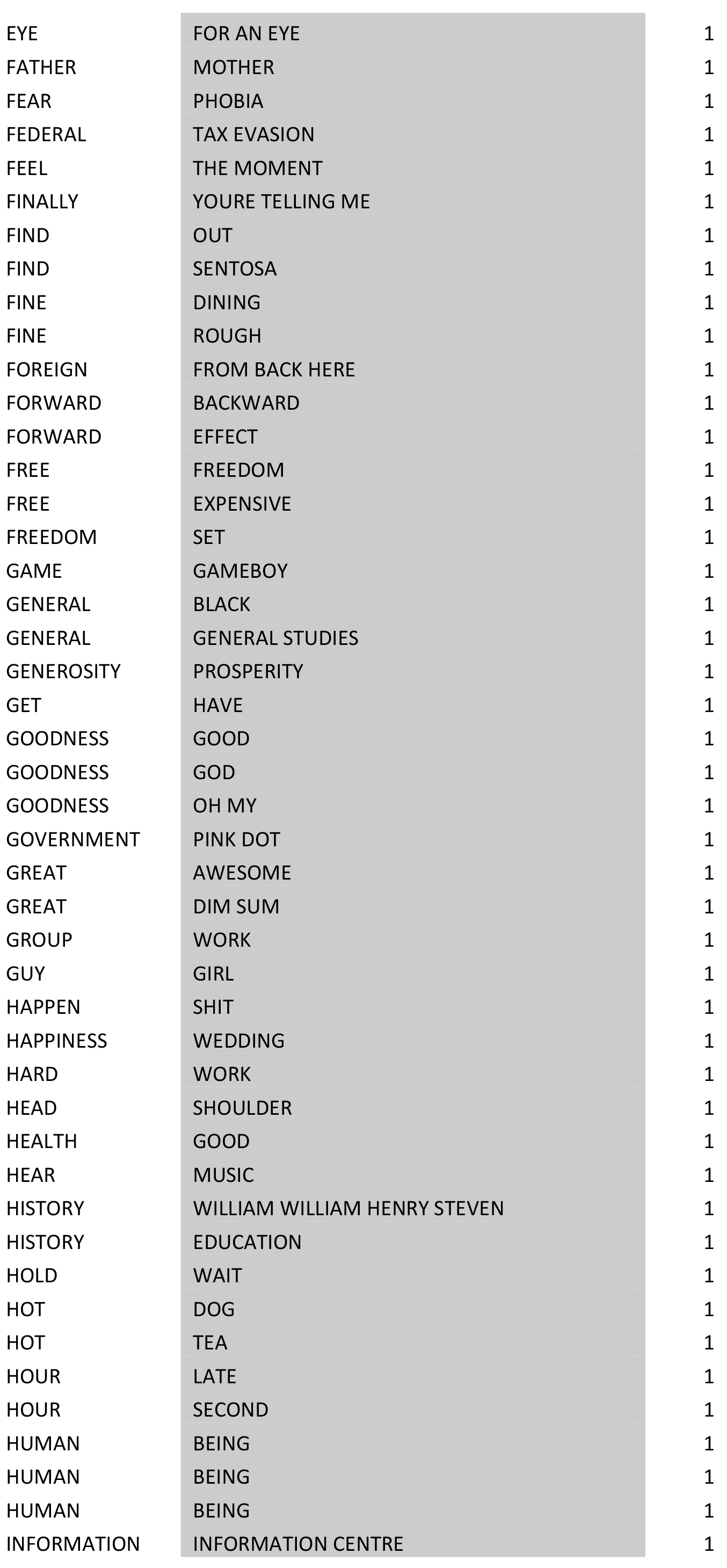

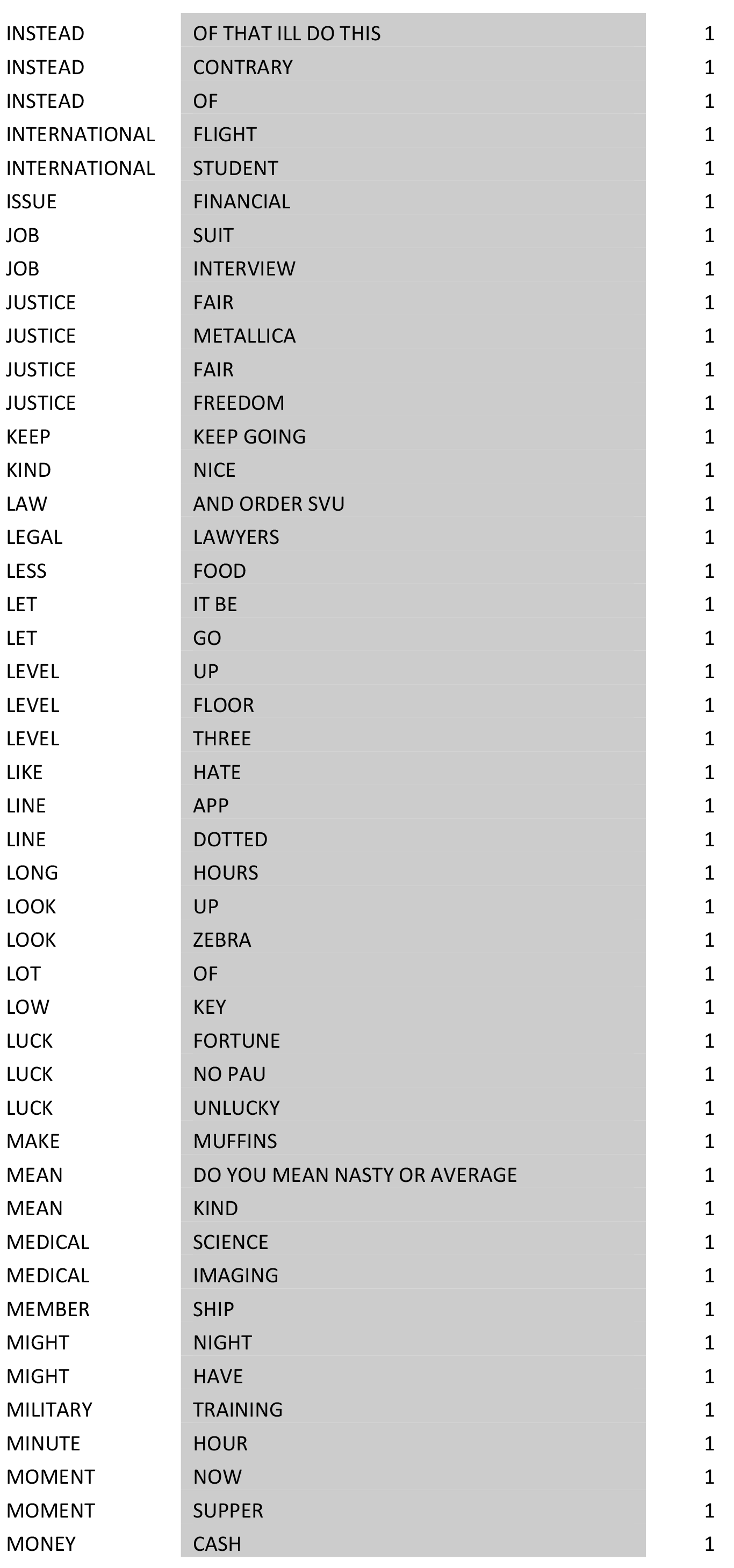

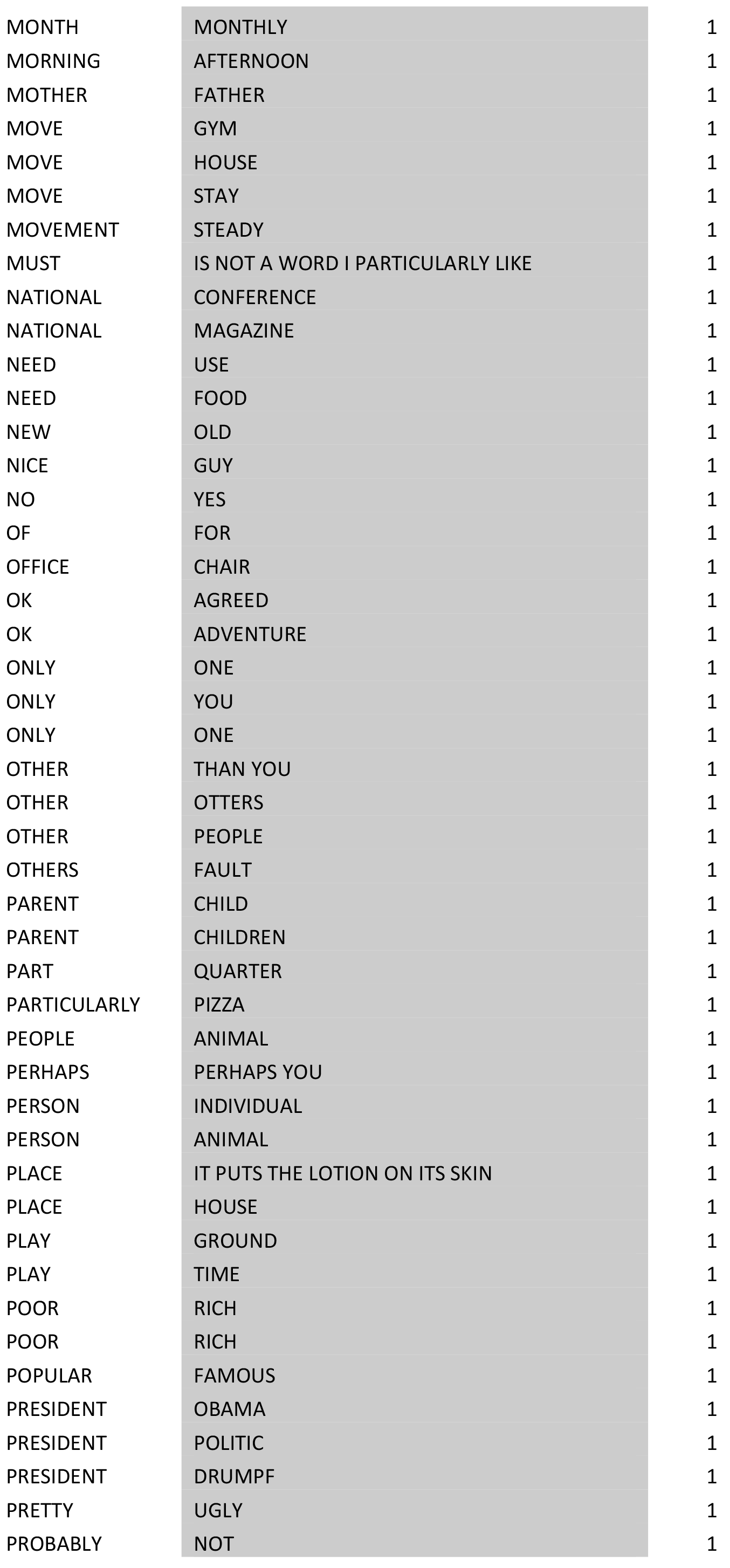

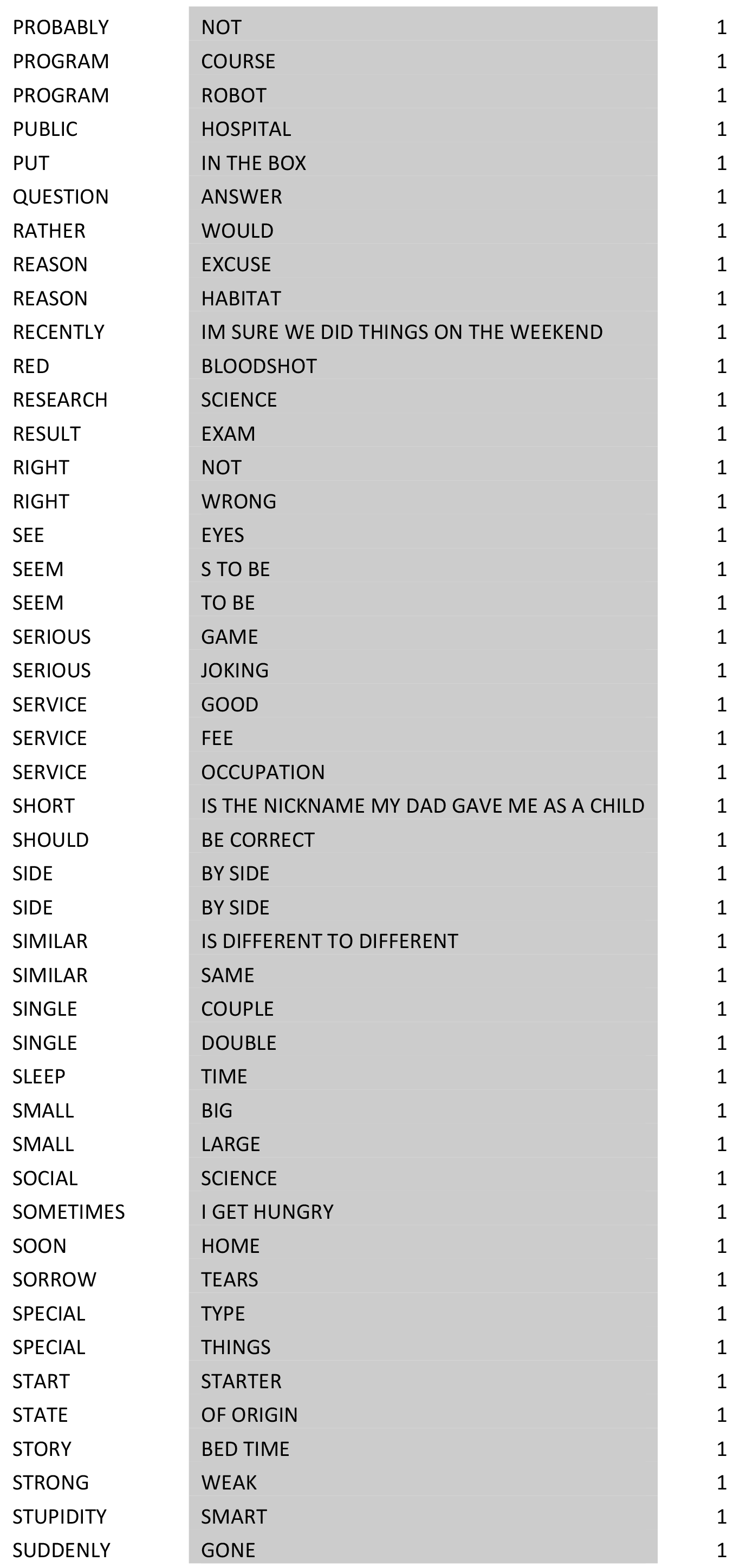

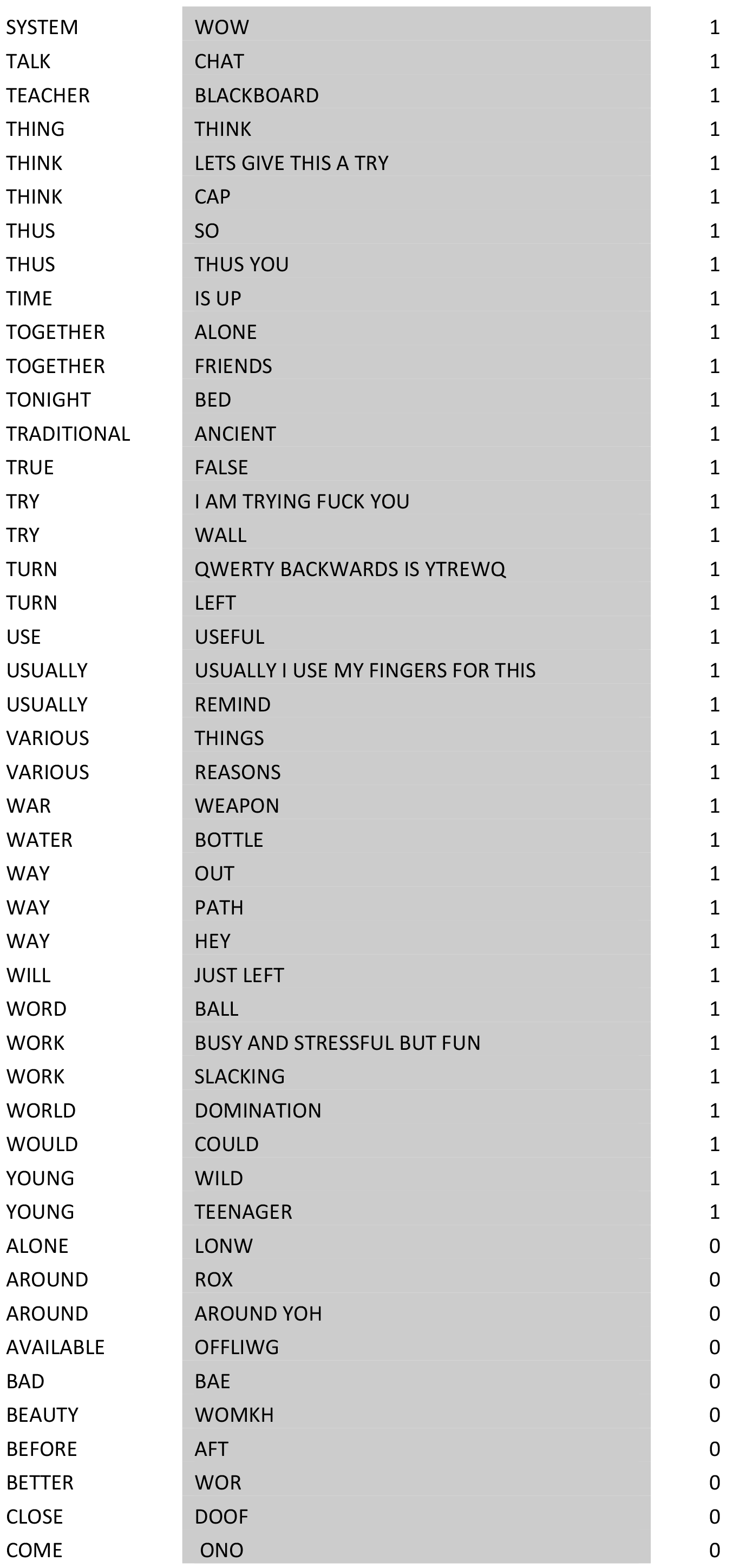

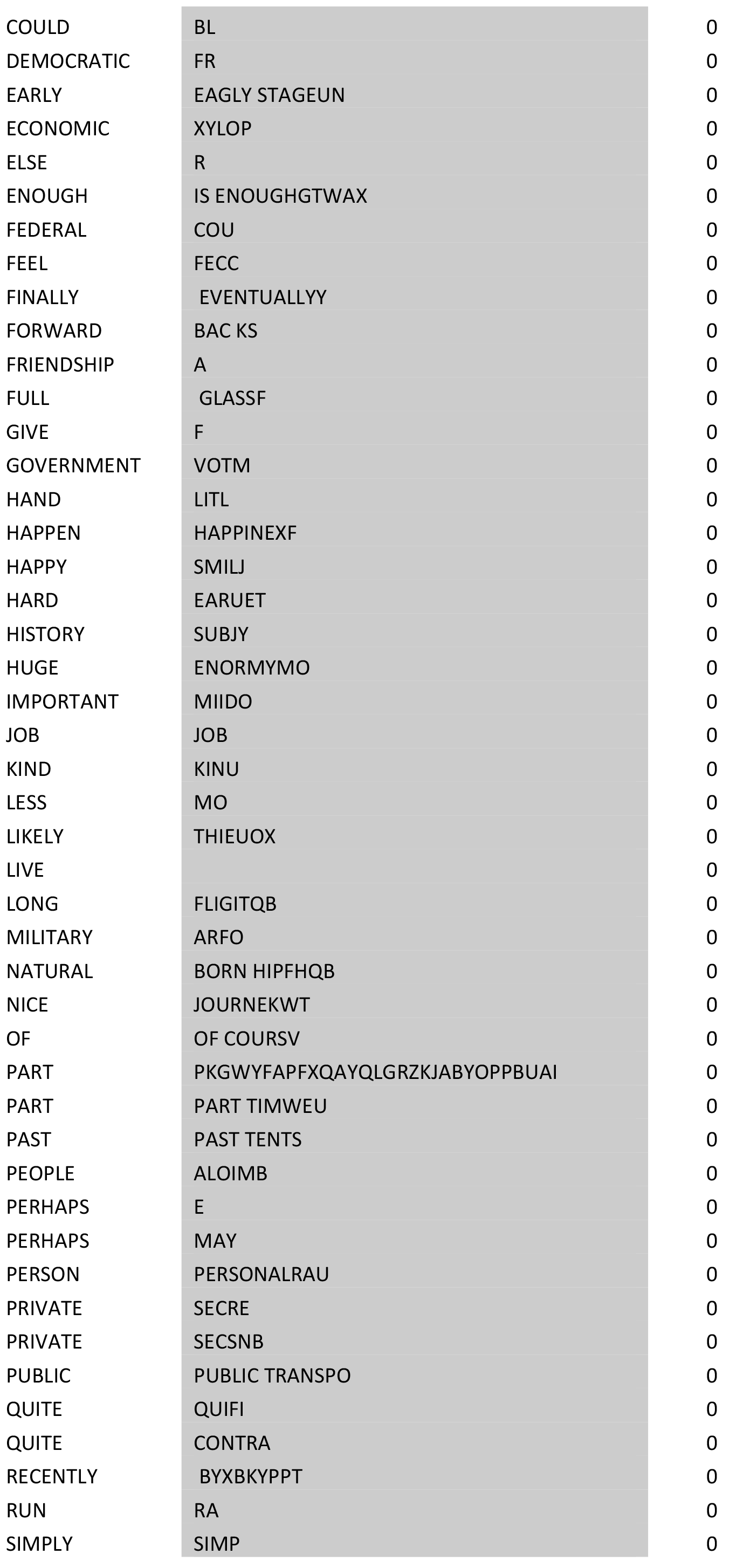

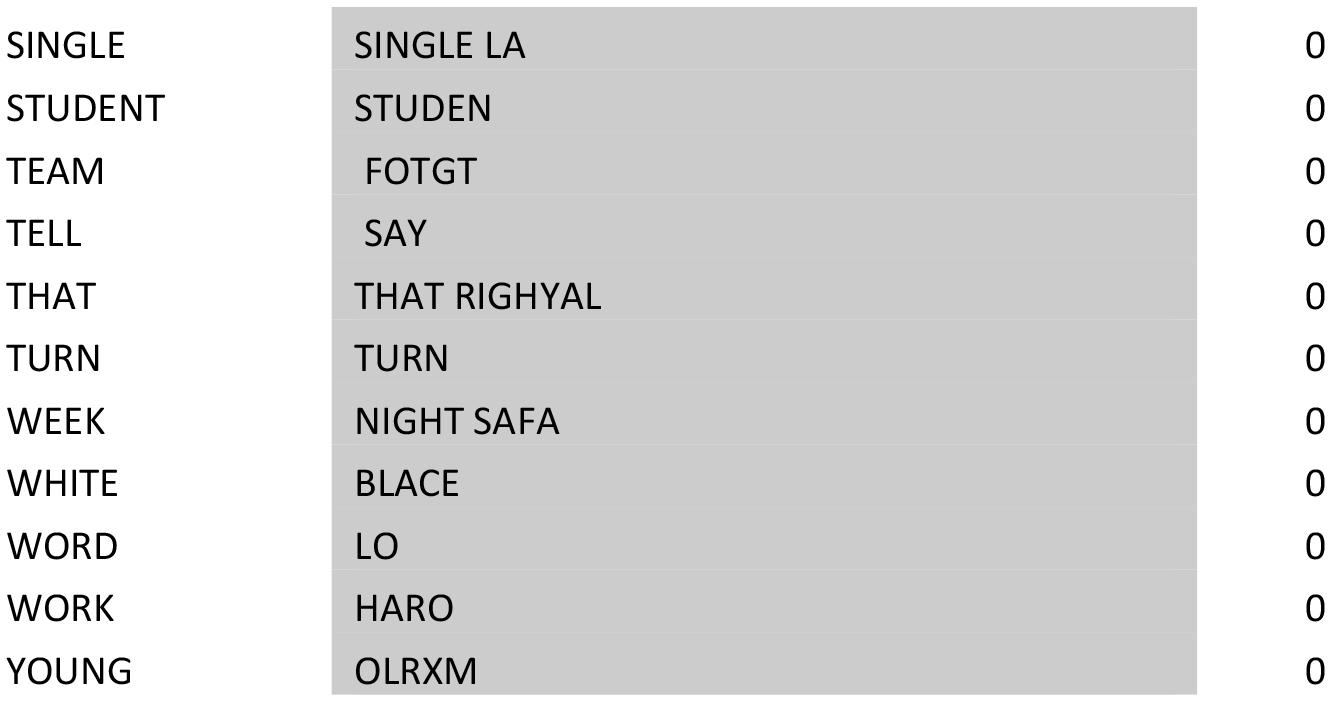

## Appendix 3. Unedited transcript of the brain-to-brain free communication of Experiment 2

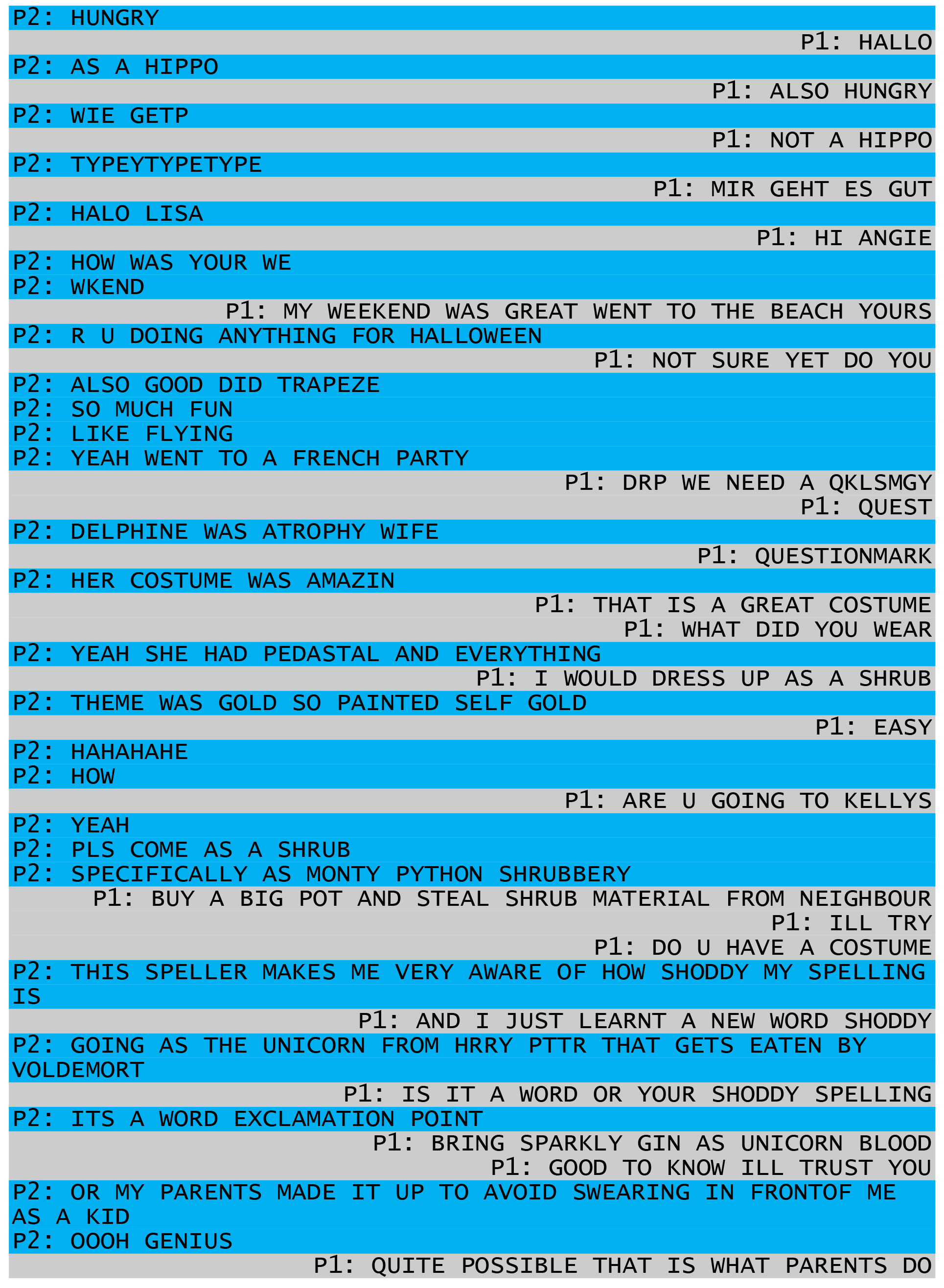

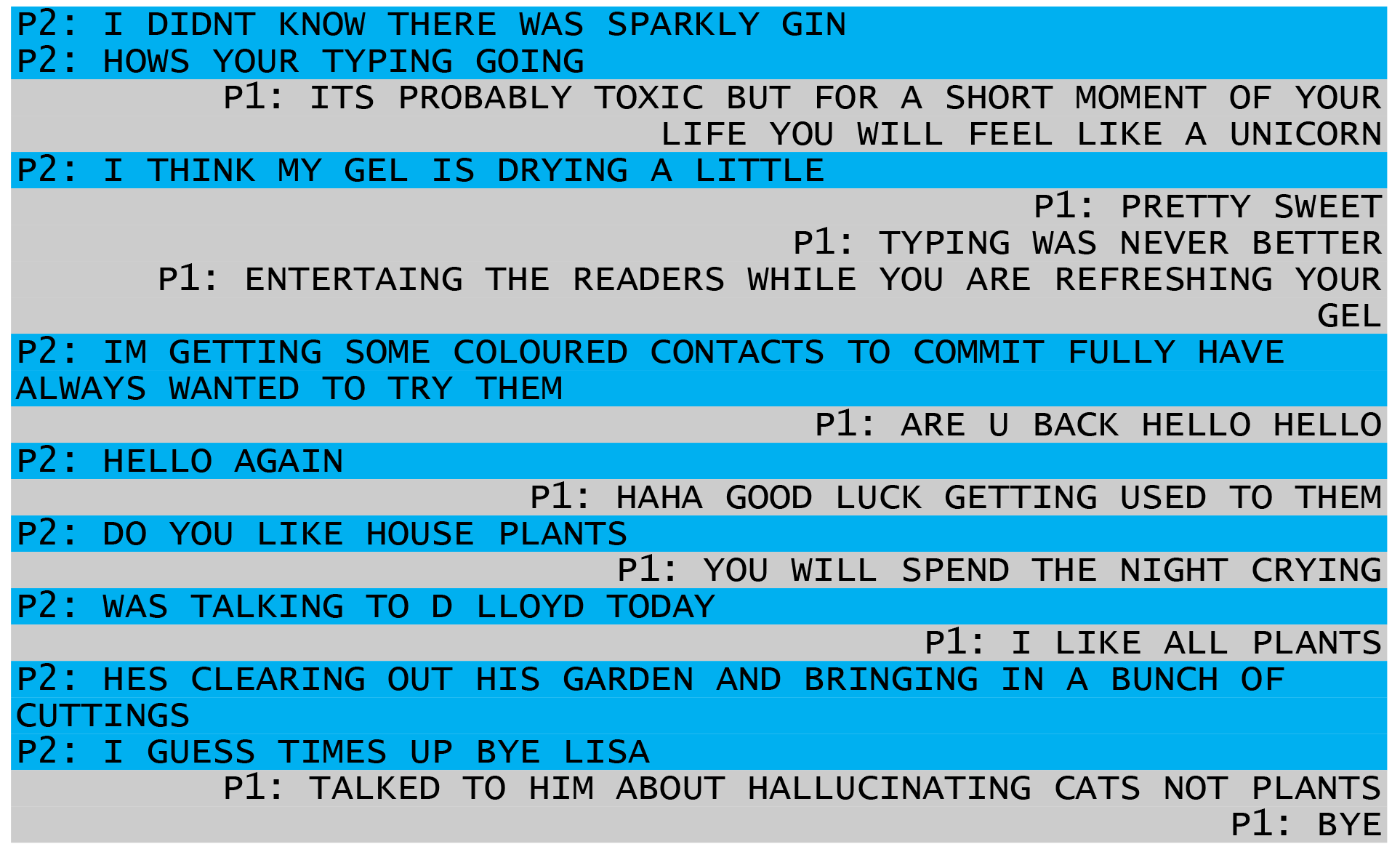

## Footnotes

* Based on Bin et al. (2011; 108 bpm), Volosyak (2011, 124 bpm), Spuller et al. (2012, 144 bpm), Chen et al. (2015, *J Neural Eng*, 151 bpm), Nakanishi et al. (2014, 167 bpm), and Sengelmann et al. (2017; 181 bmp)

